# Kinesin-1 selectivity for stable microtubules is mediated by MAP7D1’s recognition of expanded lattices

**DOI:** 10.64898/2026.11.15.766762

**Authors:** Jasper van Schelt, Varsha Mahapatra, Malina K. Iwanski, Daphne Jurriens, Niels Pagani, Anthea Meijer, Wendy van Straaten, Jitske Bak, Wilco Nijenhuis, Anna Akhmanova, Lukas C. Kapitein

## Abstract

Microtubules are organized into functionally distinct subsets that differ in stability, post-translational modifications (PTMs) and lattice spacing, but what defines these subsets and governs selective binding of motors and microtubule-associated proteins (MAPs) to specific subsets has remained unclear. Here, we demonstrate that the preferential motility of the motor protein Kinesin-1 along stable, long-lived microtubules depends on microtubule lattice spacing and MAP7 family members. Using Kinesin-1 truncations and MAP7 family depletions, we show that Kinesin-1 subset selectivity is not specified by its motor domain, but instead requires its coiled-coil 1 region, which mediates MAP7 binding. While depletion of all MAP7 proteins abolishes Kinesin-1 motility, loss of MAP7D1 alone results in non-selective motility, identifying MAP7D1 as the key determinant of selectivity. Using knock-out cells lacking detyrosinated and acetylated microtubules, we find that MAP7D1 enrichment on stable microtubule subsets depends on lattice structure rather than PTMs, whereas *in vitro* reconstitution assays directly demonstrate that MAP7D1 preferentially binds expanded microtubule lattices characteristic of long-lived microtubules. Together, our results establish microtubule lattice conformation as a key determinant of cellular microtubule subset identity and selective binding of MAPs.

## INTRODUCTION

Cell organization and intracellular transport depend on microtubules, polarized protein polymers organized into functionally distinct subsets that differ in dynamics, stability, and composition (Akhmanova & Kapitein, 2022). Dynamic microtubules grow and shrink rapidly, whereas stable microtubules are long-lived, resistant to depolymerization, and carry post-translational modifications (PTMs) such as acetylation and detyrosination (Kreis, 1987; Schulze & Kirschner, 1987). It has long been assumed that these PTMs constitute a molecular code that specifies microtubule identity and governs the differential recruitment of motors and microtubule-associated proteins (MAPs) (Janke & Magiera, 2020; McKenna et al., 2023; Verhey & Gaertig, 2007). However, whether PTMs are the major determinant of different microtubule subsets, or whether their identity relies on other microtubule features, has remained unclear.

A growing body of recent work has identified microtubule lattice conformation as an additional contributor to subset identity. While early work had shown that hydrolysis of tubulin-bound GTP upon polymerization triggers a compaction of the lattice from 8.4 to 8.2 nm (Hyman et al., 1995) and subsequent structural work has demonstrated that paclitaxel can switch GDP-bound microtubules back to the expanded conformation (Alushin et al., 2014; Vangos et al., 2026), more recent *in vitro* work revealed that several MAPs (such as Tau, doublecortin, and GEFH1) can also recognize or modulate expanded or compacted lattice conformations (Bechstedt et al., 2014; Ettinger et al., 2016; Meiring et al., 2026; Paquette et al., 2025; Siahaan et al., 2022). Tubulin-modifying enzymes have similarly been shown to exhibit sensitivity to lattice configuration (Egoldt et al., 2025; Yue et al., 2023). Notably, microtubules with distinct lattice states coexist within cells, where expanded lattices correspond to long-lived microtubules (de Jager et al., 2025). In addition, axons of iPSC-derived neurons acquire microtubules with an expanded lattice during neuronal differentiation (Zehr et al., 2026). Despite this progress, it has remained unclear whether lattice state alone, in the absence of PTMs, is sufficient to define microtubule subset identity and guide selective binding of MAPs to microtubule subsets.

Kinesin-1 provides an excellent case to address this question. This motor drives long-range intracellular transport and has long been known to move selectively along stable, highly modified microtubules in cells (Cai et al., 2009; Dunn et al., 2008; Reed et al., 2006). Nonetheless, *in vitro* reconstitution experiments with microtubules bearing different PTMs have revealed only modest effects of these modifications on Kinesin-1 motility (Dunn et al., 2008; Fernandes et al., 2026; Kaul et al., 2014; Sirajuddin et al., 2014), leaving the molecular basis for its subset selectivity unresolved. More recent work has implicated MAPs in modulating Kinesin-1 activity: Tau inhibits Kinesin-1 motility (Dixit et al., 2008; Siahaan et al., 2019; Vershinin et al., 2007), while MAP7 functions as an essential cofactor that enhances microtubule binding and processivity (Barlan et al., 2013; Chaudhary et al., 2019; Hooikaas et al., 2019; Metzger et al., 2012; Monroy et al., 2018; Sung et al., 2008; Tymanskyj et al., 2018), but limits motility at higher concentrations (Ferro et al., 2022). Intriguingly, Kinesin-1 has also been shown to expand the microtubule lattice *in vitro* (Nakata et al., 2011; Peet et al., 2018; Shima et al., 2018), raising the possibility that it could both read and remodel lattice state within cells. However, whether Kinesin-1 selectivity is mediated directly through recognition of lattice conformation or PTMs, or indirectly through MAPs that themselves recognize specific PTMs or lattice states, is unknown.

Kinesin-1 is a heterotetramer of two heavy chains, which homodimerize via coiled coils to form a central stalk, and two light chains that attach to the C-terminal part of this stalk (Yildiz, 2025). The N-terminal motor domain of the heavy chain drives microtubule-based motility, while cargo binds via the light chains and the C-terminal tail of the heavy chain. In the absence of cargo binding, kinesin adopts an autoinhibited conformation in which the tail region docks near the motor domains, facilitated by various flexible regions that interrupt the coiled coils of the heavy chain (Dietrich et al., 2008; Kaan et al., 2011; Yildiz, 2025). The motile properties of Kinesin-1 are therefore typically studied using dimeric motor constructs in which the tail region has been removed to relieve tail-mediated autoinhibition. Importantly, recent work has proposed that autoinhibition involves multiple regions and also suggested that activation is promoted by both cargo adaptors and MAP7 (Ashaduzzaman et al., 2026; Tan et al., 2023). Nonetheless, the mechanisms that ensure the subset-selective activity of Kinesin-1 have remained unresolved.

Here, we use Kinesin-1 truncation constructs, depletion of MAP7-family members, and modulation of PTMs to dissect the mechanisms underlying Kinesin-1 subset selectivity. We show that selectivity is not encoded within the Kinesin-1 motor domain but instead requires a coiled-coil region distal from its microtubule-binding site that mediates its interaction with MAP7 family members. Consistently, in COS7 cells and U2OS cells Kinesin-1 selectivity is conferred by MAP7D1, whose own enrichment on stable microtubules is governed by lattice spacing rather than PTMs. Together, our results identify microtubule lattice spacing as a key determinant of subset identity through selective recruitment of MAPs and motors.

## RESULTS

### The subset-specific microtubule binding of Kinesin-1 is specified outside its motor domain

To identify the domain(s) responsible for the microtubule subset-selectivity of Kinesin-1 (KIF5B), we started by truncating the motor to remove coiled-coil regions in its stalk, focusing on two variants that either include (K560) or exclude (K370) coiled-coil 1 (CC1) (Fig. 1A,B). We expressed these constructs fused to mStayGold in COS-7 cells together with mCherry-tubulin and performed live-cell TIRF microscopy. In agreement with previous observations (Cai et al., 2009; Reed et al., 2006), K560 moved processively along a subset of microtubules (Fig. 1C). Remarkably, K370 instead moved along all microtubules marked by mCherry-tubulin, demonstrating that Kinesin-1 subset selectivity is not specified by its motor domain but instead depends on CC1.

**Figure 1:**
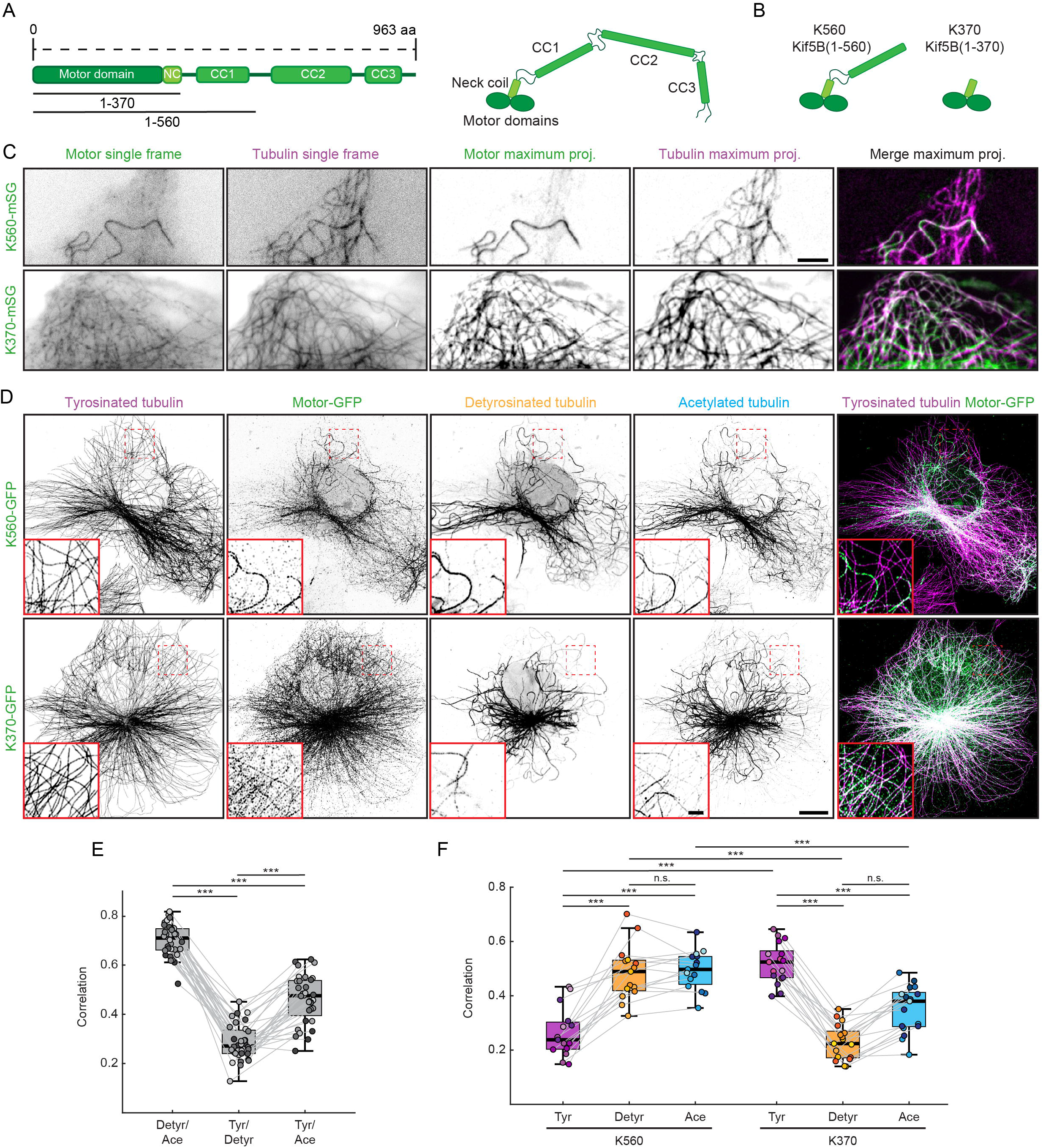
The subset-specific microtubule binding of Kinesin-1 is specified outside its motor domain. **A)** Schematic overview of the domain structure of Kinesin-1. **B)** Schematic of the Kinesin-1 truncations used in panels C-F. **C)** COS-7 cells transfected at low levels with mCherry-α-tubulin (Tubulin) and Kif5B(1-560)-mStayGold (K560-mSG) or Kif5B(1-370)-mStayGold (K370-mSG) were imaged using TIRF microscopy. Panels 1-2 show single frames while panels 3-5 show maximum projections of 20 frames, covering 2 s. Scale bar, 5 µm. **D)** COS-7 cells transfected at low levels with K560-GFP or K370-GFP were fixed using an optimized extraction-fixation protocol to visualize microtubule-bound motor before staining for tyrosinated α-tubulin (Tyr), GFP, detyrosinated α-tubulin (Detyr) and acetylated α-tubulin at position K40 (Ace). Scale bar, 10 µm (for whole cell) and 2 µm (for zoom). **E-F)** Plots showing mean ± standard deviation of Pearson’s correlation coefficient between channels of (D). 10x10µm regions were chosen from the periphery of the cell where microtubules show minimal overlap. Dots indicate individual 10 × 10 µm regions (n = 30 regions in (E) and n = 15 regions in (F) from 3 independent experiments), and shades indicate experimental repeats. A Wilcoxon matched-pairs signed rank test was used to test for statistical significance between correlations of the same regions (E and within condition in F) and a Welch’s t-test was used to compare between conditions (K560 vs K370 in F). ns = not significant, *** = p<0.001.

To further analyze this finding, we next explored the localization of K560 and K370 in fixed cells stained for acetylated, tyrosinated and detyrosinated tubulin. We optimized an extraction-fixation protocol that removes the unbound fraction of active motor to visualize the microtubule-bound fraction together with these different microtubule subsets. In addition, the signal of GFP-tagged constructs was boosted with an anti-GFP antibody, allowing us to minimize overexpression artifacts by analyzing the lowest expressing cells. As expected, detyrosination and acetylation mostly marked the same microtubules in COS-7 (Fig. 1D) and U2OS cells (Fig. S1A) and hence showed a high correlation coefficient (Fig. 1E). In contrast, tyrosination marked a different microtubule population (Fig. 1D) and therefore had a low correlation coefficient with detyrosination and acetylation (Fig. 1E). Overexpression of the motor constructs revealed that K560 was strongly enriched on detyrosinated and acetylated compared to tyrosinated microtubules (Fig. 1D,F, Fig. S1A). In contrast, K370 bound to all microtubules and showed an increased correlation with tyrosinated microtubules and a decreased correlation with detyrosinated and acetylated microtubules (Fig. 1D,F, Fig. S1A). To exclude the possibility that loss of selectivity resulted from inefficient dimerization of K370, we introduced a GCN4 leucine zipper to enforce dimerization. K370-GCN4 similarly bound to all microtubule subsets, indicating that loss of selectivity was not caused by impaired dimerization (Fig. S1B). Together, these data demonstrate that the selectivity of Kinesin-1 for detyrosinated and acetylated microtubules is not exclusively encoded in the motor domain but instead depends on the presence of the CC1 region.

### Differential microtubule subset recognition by Kinesin-1 activating MAP7-family members

Previous *in vitro* studies have shown that binding of MAP7-family proteins to residues 500-520 within the Kinesin-1 CC1 region increases the landing rate and run length of Kinesin-1, presumably by relieving a form of autoinhibition imposed by the CC1 region (Hooikaas et al., 2019; Monroy et al., 2018). All MAP7-family members (MAP7, MAP7D1, MAP7D2 and MAP7D3) can interact with Kinesin-1 through a conserved C-terminal region, as well as with microtubules through a conserved N-terminal region (Fig. 2A). We therefore hypothesized that the subset selectivity of Kinesin-1 was related to the localization of MAP7-family members to specific microtubule subsets (Masson & Kreis, 1993). To test this, we first determined the expression levels of MAP7 proteins in commonly used cell lines by Western blot (Fig. 2B). MAP7D2 expression was not detected in any of the tested cell lines, in agreement with its neuron-specific function (Pan et al., 2019). In contrast, MAP7D3 was robustly expressed in all cell lines examined. MAP7 and MAP7D1 were expressed to varying degrees, with MAP7 showing higher expression in HeLa cells and MAP7D1 higher expression in U2OS cells (Fig. 2B). To determine the subset-specific localization of MAP7-family members, we expressed GFP-fusion proteins in COS-7 cells at low levels and stained for microtubule modifications (Fig. 2C). MAP7D1 and MAP7D2 showed high specificity for acetylated and detyrosinated microtubules. MAP7 was less selective compared to MAP7D1 and MAP7D2 because it also showed low signal on tyrosinated microtubules towards the center of the cell (Fig. 2C). In contrast, MAP7D3 was mostly diffusely localized, and only enriched on microtubules in highly expressing cells. Staining for endogenous proteins in COS-7 and U2OS cells confirmed that MAP7D1 is specifically enriched on acetylated microtubules while MAP7D3 is mostly diffuse with some enrichment near the microtubule organizing center (Fig. 2D, Fig. S2A).

**Figure 2:**
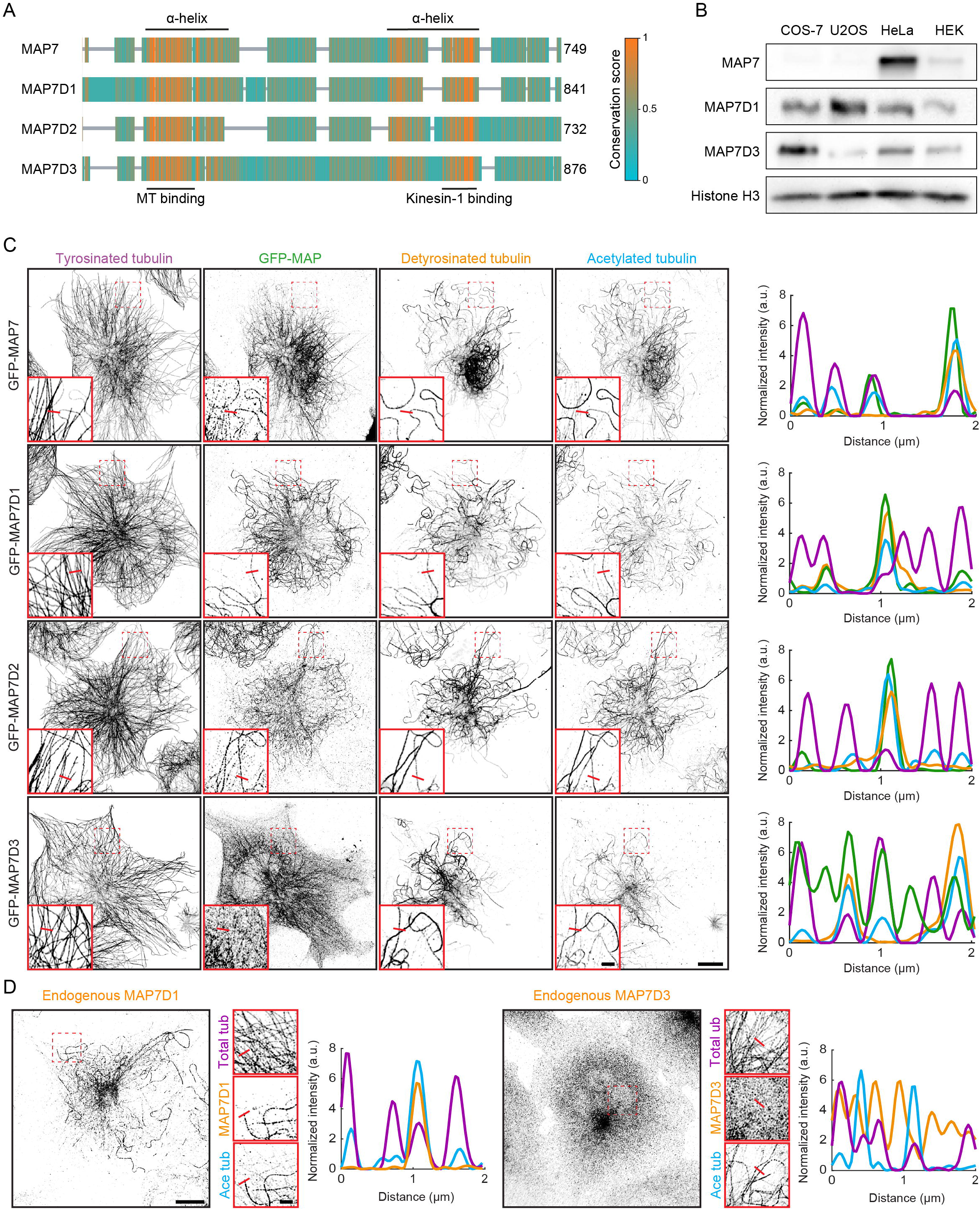
MAP7-family members differ in expression between cell types and in microtubule subset binding. **A)** Alignment and sequence conservation of human MAP7-family proteins MAP7, MAP7D1, MAP7D2 and MAP7D3. All MAP7-family proteins contain a conserved N-terminal microtubule binding domain and a conserved C-terminal Kinesin-1 binding domain. **B)** Western blot of the expression of MAP7-family proteins in extracts of COS-7, U2OS, HeLa and HEK293T (HEK) cells. Histone H3 was used as a loading control. **C)** COS-7 cells transfected at low levels with GFP-tagged MAP7-family proteins were fixed before staining for tyrosinated α-tubulin, GFP, detyrosinated α-tubulin and acetylated α-tubulin at position K40. Red lines in the zooms of C and D indicate 2 µm long regions used for the corresponding intensity plots. **D)** COS-7 cells were fixed and stained for α-tubulin and β-tubulin (Total tub), acetylated α-tubulin at position K40 (Ace tub) and endogenous MAP7D1 or MAP7D3. Scale bars, 10 µm (C,D whole cell) and 2 µm (C,D zoom).

Finally, to determine whether MAP7D1-positive microtubules correspond to the stable microtubule population, we generated a U2OS MAP7D1-Halo knock-in cell line and followed microtubule dynamics by live-cell TIRF microscopy (Fig. S2B). Upon nocodazole treatment, most microtubules depolymerized, whereas MAP7D1-positive microtubules persisted. Following nocodazole washout, newly polymerized microtubules were initially MAP7D1-negative, further distinguishing these dynamic microtubules from the pre-existing MAP7D1-positive population. Together, these observations demonstrate that MAP7-family members differentially associate with distinct microtubule populations, with MAP7D1 selectively localizing to the subset of stable microtubules.

### MAP7D1 is required for Kinesin-1’s microtubule subset selectivity

To directly examine the relation between the selectivity of Kinesin-1 and MAP7D1, we first expressed K560-GFP in COS7 cells and stained for endogenous MAP7D1. K560 largely colocalized with endogenous MAP7D1, specifically on acetylated microtubules (Fig. 3A). Consistent with this, live-cell imaging in U2OS MAP7D1-Halo knock-in cells showed that K560 preferentially moved along MAP7D1-positive microtubules, whereas K370 did not have this preference and also moved along MAP7D1-negative microtubules (Fig. S3A). Next, we explored the contribution of MAP7-family proteins to Kinesin-1 selectivity using siRNA-mediated knockdown of endogenous MAP7D1 and MAP7D3 in COS-7 cells. Western blot analysis confirmed efficient knockdown with no upregulation of other MAP7-family proteins (Fig. S3B). After siRNA transfection we expressed K560-GFP, followed by fixation and staining for microtubule modifications to visualize different microtubule subsets. Depletion of MAP7D1 reduced colocalization of K560 with detyrosinated and acetylated microtubules compared to controls (Fig. 3B, Fig. S3C). Interestingly, the colocalization with tyrosinated microtubules increased, suggesting that K560 is no longer subset-selective upon loss of MAP7D1. Depletion of MAP7D3, on the other hand, did not significantly affect the localization of K560. Simultaneous depletion of MAP7D1 and MAP7D3 caused diffuse localization of K560 and low correlation with all microtubule subsets, indicating that K560’s ability to bind microtubules was strongly impaired. We confirmed these findings in U2OS cells (Fig. S3D). Importantly, simultaneous knockdown of MAP7D1 and MAP7D3 did not affect the microtubule binding of K370, indicating that the reduced microtubule association observed for K560 depends on the presence of the CC1 domain (Fig. 3B, Fig. S3C). Together, these data indicate that, inside cells, binding of diffusely located MAP7D3 to CC1 drives generic activation of Kinesin-1 and motility on all subsets, whereas the localization of MAP7D1 to detyrosinated and acetylated microtubules restricts activation and motility of Kinesin-1 to this specific subset. Furthermore, Kinesin-1’s selectivity in wildtype conditions suggests that MAP7D1 is the dominant activator in these conditions.

**Figure 3:**
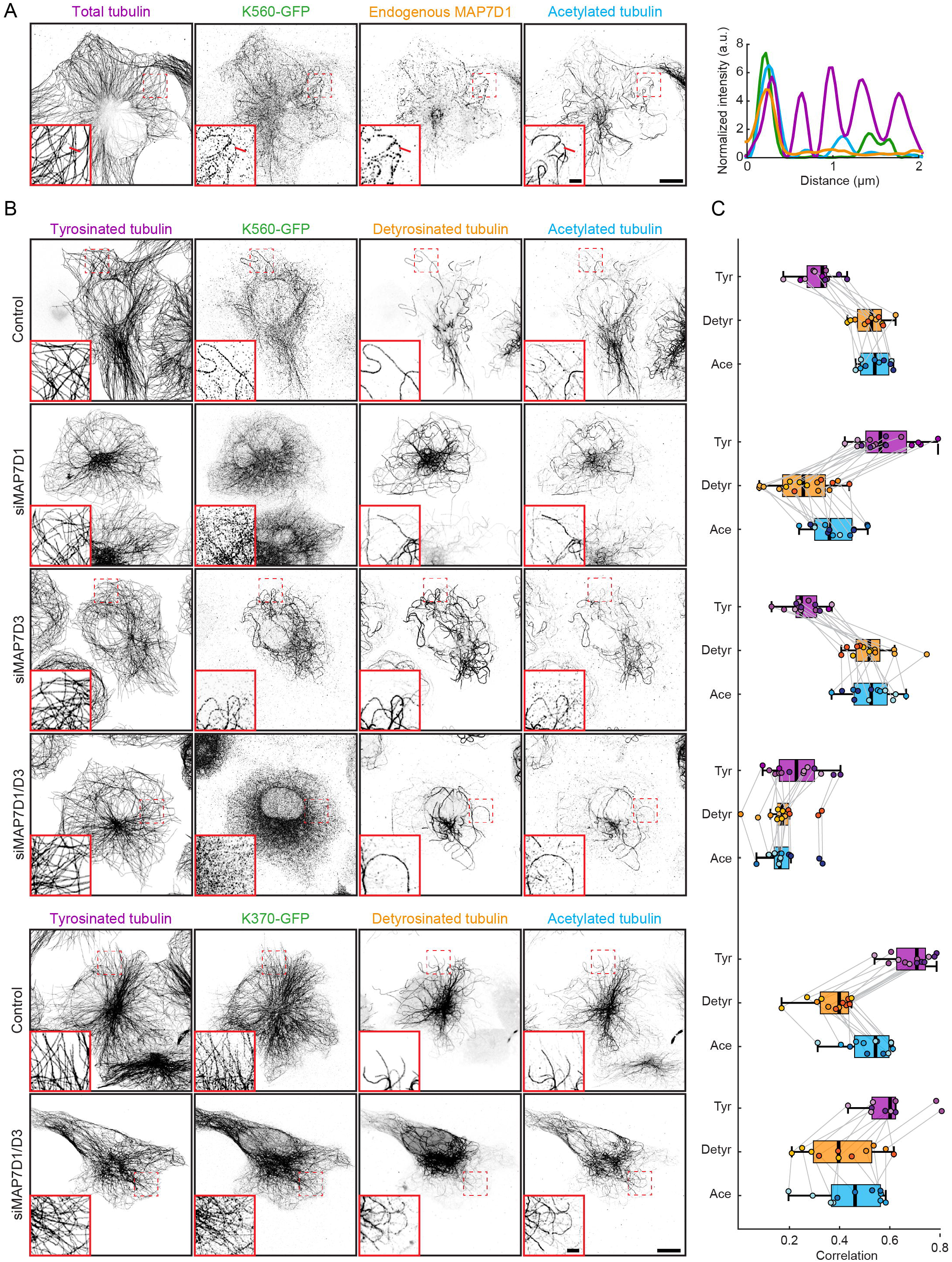
MAP7D1 is required for Kinesin-1’s microtubule subset selectivity in COS-7 cells. **A)** COS-7 cells transfected at low levels with Kif5B(1-560)-GFP (K560-GFP) were fixed and stained for α-tubulin and β-tubulin (Total tubulin), GFP, endogenous MAP7D1 and acetylated α-tubulin at position K40. Red lines in the zooms indicate 2 µm long regions used for the corresponding intensity plots. **B)** COS-7 cells transfected with siRNAs for Luciferase (Control), MAP7D1, MAP7D3 or double MAP7D1/MAP7D3 before overexpression of K560-GFP or K370-GFP at low levels. Cells were fixed and stained for tyrosinated α-tubulin (Tyr), GFP, detyrosinated α-tubulin (Detyr) and acetylated α-tubulin at position K40 (Ace). Scale bars, 10 µm (whole cell) and 2 µm (zoom). **C)** Plots showing Pearson’s correlation coefficients between channels in B, calculated from 10 x 10µm regions from the periphery of the cell where microtubules show minimal overlap. Dots indicate individual 10 × 10 µm regions (n = 16 regions from 3 independent experiments), and shades indicate experimental repeats. A Wilcoxon matched-pairs signed rank test was used to test for statistical significance between subset correlations of the same condition and a Welch’s t-test was used to compare between conditions. Significance is shown in Fig. S3C.

### Detyrosination, but not acetylation, increases microtubule binding of MAP7D1

Given that MAP7D1 is required for Kinesin-1 selectivity and is highly enriched on detyrosinated and acetylated microtubules, we next investigated the mechanism of MAP7D1 subset recognition. To this end, we treated GFP-MAP7D1 expressing COS-7 cells with the drug tubacin, which inhibits the deacetylase HDAC6 and increases acetylation on all microtubules (Fig. 4A). Despite global hyperacetylation, MAP7D1 remained restricted to detyrosinated microtubules, indicating that acetylation does not determine MAP7D1 subset selectivity. Consistently, K560-GFP remained enriched on a subset of microtubules following tubacin treatment (Fig. S4A). Next, to increase the overall levels of detyrosination we overexpressed the detyrosinating enzyme VASH1 and its cofactor SVBP (Aillaud et al., 2017; Nieuwenhuis et al., 2017), in combination with GFP-MAP7D1 (Fig. 4A). As expected, this resulted in increased detyrosination and decreased tyrosination compared to non-transfected cells. In these conditions, MAP7D1 remained enriched on a subset of microtubules, although it was now also present to some extent on the more straight and radially oriented microtubules.

**Figure 4:**
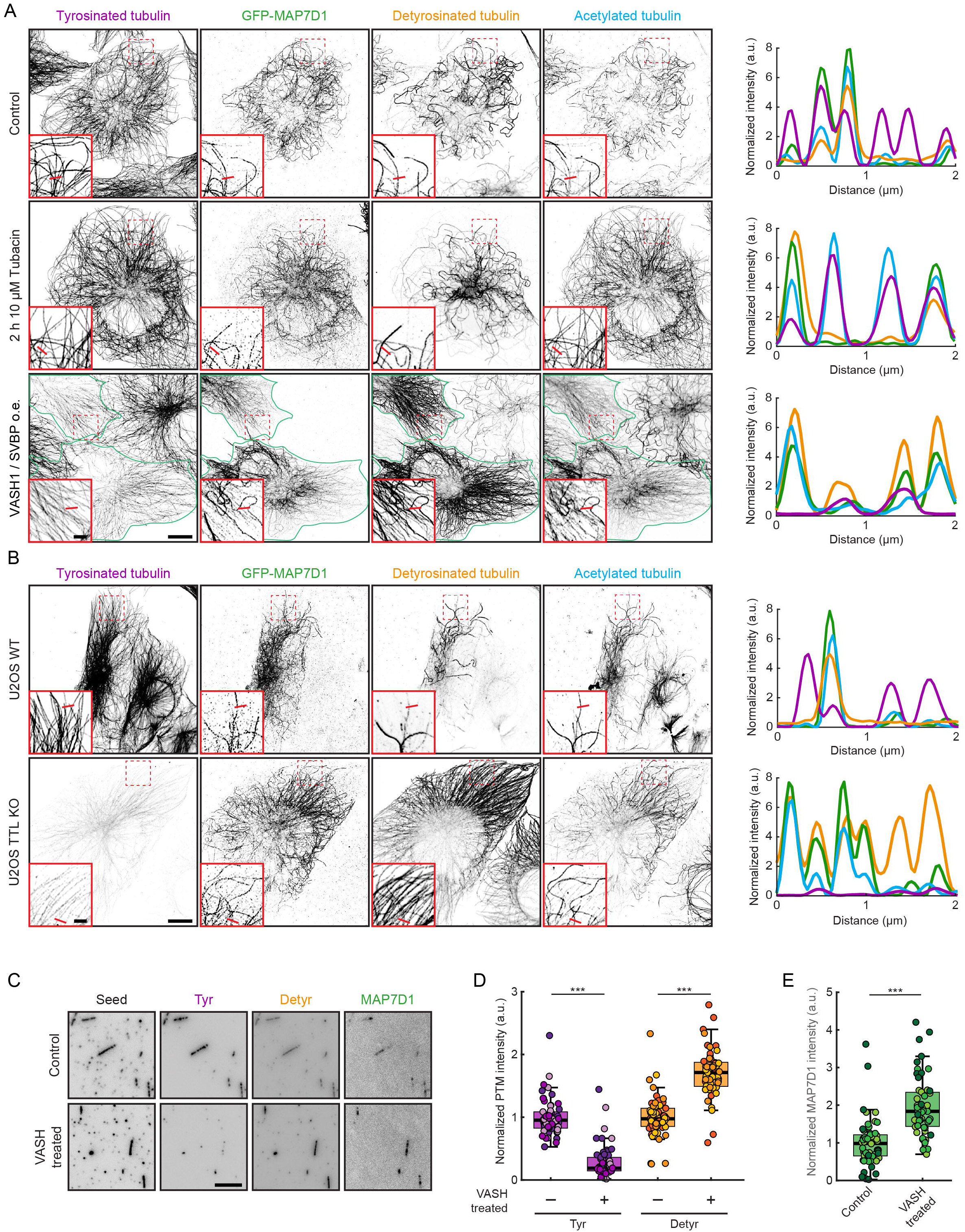
Microtubule subset selectivity of MAP7D1 is partially mediated by detyrosination and not by acetylation. **A)** COS-7 cells transfected at low levels with GFP-MAP7D1 were treated with DMSO (control), 10µM Tubacin for 2 h or co-transfected with VASH1-FLAG/SVBP-FLAG before fixing and staining for tyrosinated α-tubulin, GFP, detyrosinated α-tubulin and acetylated α-tubulin at position K40. Green outlines in VASH1/SVBP indicate cells co-transfected with GFP-MAP7D1 and VASH1-FLAG/SVBP-FLAG. Red lines in the zooms of A and B indicate 2 µm long regions used for the corresponding intensity plots. **B)** Wildtype and TTL KO U2OS cells transfected at low levels with GFP-MAP7D1 were fixed and stained for tyrosinated α-tubulin, GFP, detyrosinated α-tubulin and acetylated α-tubulin at position K40. **C)** GMPCPP microtubule seeds were incubated *in vitro* with MRB80 buffer (Control) or with MRB80 buffer containing purified VASH1 and SVBP (VASH treated) for 45 min followed by incubation with purified fluorescently labeled MAP7D1 for 10 min. Seeds were subsequently fixed and stained for tyrosinated α-tubulin (Tyr) and detyrosinated α-tubulin (Detyr) and imaged using TIRF microscopy. **D)** Analysis of (C) showing mean ± standard deviation of Tyr or Detyr intensity normalized to control (control n = 58, treated n = 55; from 3 experiments). **E)** Analysis of (C) showing mean ± standard deviation of MAP7D1 intensity normalized to control (control n = 58, treated n = 55; from 3 experiments). A Welch’s t-test was used to compare between VASH treated and control seeds in (D,E), *** = p<0.001. Scale bars, 10 µm (A,B whole cell), 2 µm (A,B zoom), 5 µm (C).

We noted that the combined overexpression of VASH1, SVBP and GFP-MAP7D1 also impacted acetylation, which was now more evenly distributed across all microtubules compared to controls. To increase detyrosination without possible overexpression artifacts, we performed CRISPR-Cas9-mediated knockout in U2OS cells for the gene encoding the tyrosination enzyme TTL. In TTL-KO cells, the levels of tyrosination were strongly decreased while detyrosination levels were high on all microtubules (Fig. 4B). In these conditions, acetylation was mostly restricted to the subset of highly curved microtubules, similar to control cells (Fig. S4C). GFP-MAP7D1 was mostly restricted to this acetylated subset but with a mild redistribution to the more straight and radially oriented microtubules, which was also observed for endogenous MAP7D1 (Fig. 4B, Fig. S4C). Upon TTL knockout, K560 was slightly redistributed to the radial microtubules and often enriched at the periphery of the cells, consistent with increased Kinesin-1 activation on the outward oriented dynamic microtubules (Fig. S4D).

To test if detyrosination directly affects MAP7D1 binding to microtubules, we performed *in vitro* assays using purified proteins in which GMPCPP-stabilized microtubules were treated with purified VASH1 and SVBP before washing in purified MAP7D1. VASH1/SVBP treated microtubules had decreased tyrosination and increased detyrosination levels, confirming the activity of the enzymes (Fig. 4C,D). In addition, MAP7D1 binding was about twofold increased on VASH1/SVBP-treated microtubules (Fig. 4E). Together, these data demonstrate that detyrosination, but not acetylation, promotes MAP7D1 binding to microtubules but cannot fully account for its enrichment on the subset of stable microtubules.

### MAP7D1 recognizes microtubules independently of acetylation and detyrosination

Since GFP-MAP7D1 still strongly associated with a subset of microtubules in TTL-KO cells, we hypothesized that another feature of microtubules might be the primary cause of MAP7D1 selectivity. Previous work revealed that stable and highly modified microtubules have an expanded lattice in cells, whereas dynamic microtubules are compacted (de Jager et al., 2025; Jansen et al., 2023). To examine how lattice expansion affects MAP7D1 localization, we first treated GFP-MAP7D1 expressing cells with the lattice expanding drug paclitaxel for 20 minutes, a treatment previously reported to alter MAP7D1 localization (Khushalani et al., 2025). Upon treatment with paclitaxel, MAP7D1 redistributed to all microtubules (Fig. 5A, Fig. S5A). In contrast, treatment with discodermolide, a drug that stabilizes microtubules but does not expand the lattice (Meiring et al., 2026), did not result in the redistribution of MAP7D1, suggesting that lattice expansion and not stabilization controls MAP7D1 binding.

**Figure 5:**
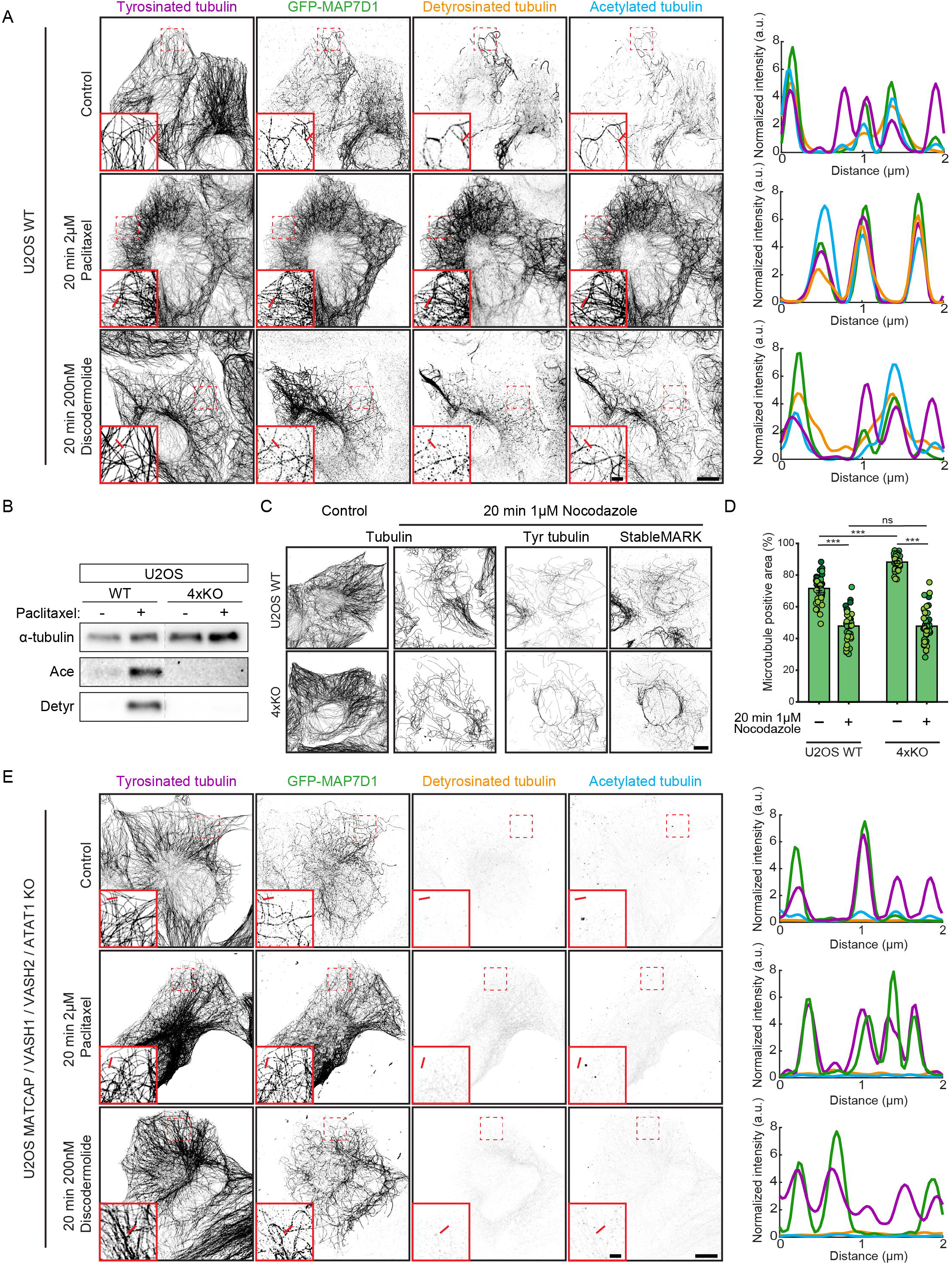
MAP7D1 recognizes microtubules with an expanded lattice independently of acetylation and detyrosination. A) Wildtype U2OS cells transfected at low levels with GFP-MAP7D1 were treated with DMSO (Control), 2 µM paclitaxel or 200 nM discodermolide for 20 min before fixation and staining for tyrosinated α-tubulin, GFP, detyrosinated α-tubulin and acetylated α-tubulin at position K40. Red lines in the zooms indicate 2 µm long regions used for the corresponding intensity plots. B) Western blot of wildtype U2OS and MATCAP/VASH1/VASH2/ATAT1 quadruple knockout (4xKO) U2OS cell extracts stained for α-tubulin, detyrosinated α-tubulin and acetylated α-tubulin at position K40 (Ace). Cells were either untreated or treated with 2µM paclitaxel prior to lysis. C) Wildtype and 4xKO U2OS cells were treated with DMSO (Control) or 1 µM nocodazole for 20 min before fixation and staining for α-tubulin and β-tubulin (Total tubulin). In separate experiments, cells expressing low levels of StableMARK-GFP were subjected to the same treatment before fixation and staining for tyrosinated α-tubulin (Tyr) and GFP. D) Quantification of nocodazole-resistant microtubules in (C), based on lower-magnification images as shown in Fig. S5A. The percentage of cell area occupied by total tubulin signal was measured. Dots indicate individual cells and shades indicate experimental repeats (n = 39 cells from 3 independent experiments). A Welch’s t-test was used to compare between conditions. ns = not significant, *** = p < 0.001. **E)** 4xKO U2OS cells transfected at low levels with GFP-MAP7D1 were treated with DMSO (Control), 2 µM paclitaxel or 200 nM discodermolide for 20 min before fixation and staining for tyrosinated α-tubulin, GFP, detyrosinated α-tubulin and acetylated α-tubulin at position K40. Red lines in the zooms indicate 2 µm long regions used for the corresponding intensity plots. Scale bars, 10 µm (whole cell) and 2 µm (zoom).

Treatment with paclitaxel also increased the levels of detyrosination and acetylation, consistent with the preference of microtubule modifying enzymes for microtubules with an expanded lattice (Egoldt et al., 2026; Yue et al., 2023). To investigate whether the increased binding of MAP7D1 upon paclitaxel treatment was directly caused by lattice expansion or by the increased levels of PTMs, we generated a U2OS quadruple knockout cell line for the detyrosinating enzymes VASH1, VASH2, and MATCAP (Landskron et al., 2022), and the acetylating enzyme ATAT1 (Akella et al., 2010; Shida et al., 2010) (U2OS-4xKO). Western blot and immunocytochemistry confirmed the complete loss of both detyrosination and acetylation (Fig. 5B, Fig. S5A-C), while cell viability and growth appeared relatively unchanged. Interestingly, StableMARK (a marker for stable microtubules that binds expanded lattices (de Jager et al., 2025; Jansen et al., 2023)) remained localized to a subset of microtubules in U2OS-4xKO cells, suggesting that these cells still feature a subset of stable microtubules (Fig. S5D). Consistent with this, we observed a persistent subset of microtubules following treatment with the microtubule destabilization agent nocodazole, indicating that detyrosination and acetylation are not required for the formation of stable microtubules (Fig. 5C,D, Fig. S5A,B). This nocodazole-resistant subset was also decorated by StableMARK at low levels of overexpression, suggesting that the remaining microtubules have an expanded lattice (Fig. 5C). Furthermore, GFP-MAP7D1 and K560 were also still localized to a subset of microtubules in the absence of acetylation and detyrosination (Fig. 5E, Fig. S5E). To further examine whether lattice conformation determines MAP7D1 localization, we next compared the effects of paclitaxel and discodermolide in the U2OS-4xKO cells. Treatment with paclitaxel, but not discodermolide, resulted in the redistribution of MAP7D1 to all microtubules, indicating that lattice expansion promotes MAP7D1 binding independently of detyrosination and acetylation (Fig. 5E).

### MAP7D1 recognizes microtubules with an expanded lattice *in vitro*

To directly test whether lattice expansion controls the binding of MAP7D1 to microtubules, we turned to an *in vitro* reconstitution assay in which we could directly trigger and observe microtubule lattice expansion (Meiring et al., 2026) (Fig. 6A). Sparsely labeled dynamic microtubules were grown from GMPCPP-stabilized seeds before wash-in of either paclitaxel or discodermolide. Whereas paclitaxel triggered a ∼2.7% increase in microtubule length, consistent with the reported increase in dimer spacing from 8.2 to 8.4 nm (Alushin et al., 2014), discodermolide did not result in lattice expansion (Fig. 6A,B). To test the effect of lattice expansion on MAP7D1 binding, we next repeated these experiments in the presence of low concentrations (500 pM) of MAP7D1. Wash-in of paclitaxel to trigger lattice expansion caused a nearly 30-fold increase in MAP7D1 intensity on microtubules (Fig. 6C,D), compared to a modest 1.3-fold increase upon wash-in of discodermolide. Together, these data demonstrate that MAP7D1 directly recognizes the expanded microtubule lattice, establishing lattice expansion as the principal determinant of MAP7D1 subset selectivity.

**Figure 6:**
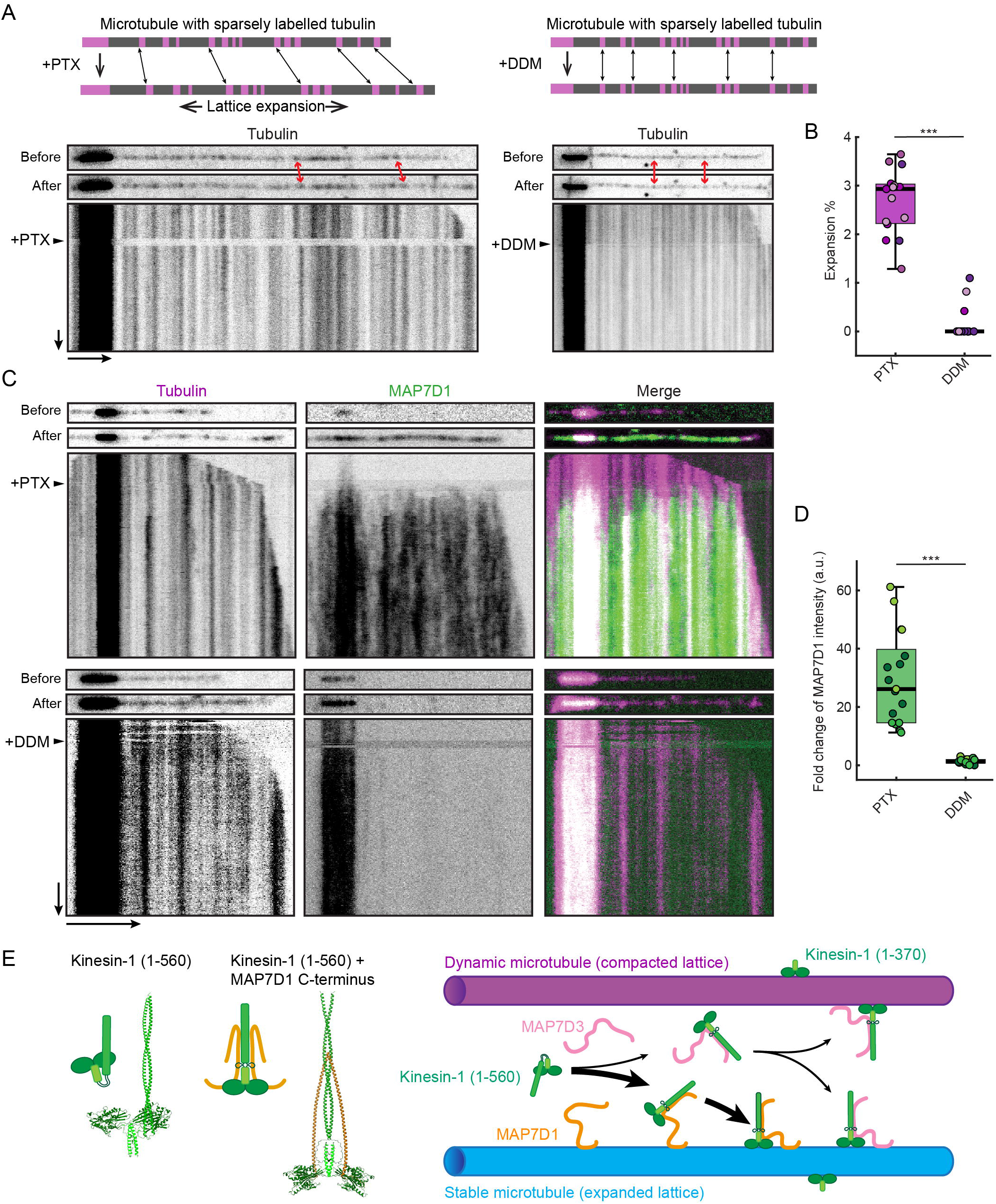
MAP7D1 recognizes microtubules with an expanded lattice *in vitro*. **A)** Top: Schematic of GMPCPP microtubule seeds with free-growing sparsely labelled dynamic ends used to detect microtubule expansion (or lack thereof) upon wash-in of paclitaxel (PTX) or discodermolide (DDM). Middle: Still images of representative microtubules before and after wash-ins, double-headed arrows highlight speckle shift upon the addition of PTX but not DDM. Bottom: Corresponding kymographs with the frame of the wash-in indicated with an arrowhead. Scale bar, 5 µm (Horizontal arrow) and 50 s (Vertical arrow). **B)** Plot from (A) showing mean ± standard deviation of percentage expansion with PTX or DDM (PTX n = 15 microtubules, DDM n = 15 microtubules; from 3 experiments). A Welch’s t-test was used to compare between PTX and DDM, *** = p<0.001. **C)** Still images of representative microtubules before and after PTX or DDM wash-in in assays with microtubules grown in the presence of 500pM MAP7D1 with corresponding kymographs with wash-in frame indicated with an arrowhead. Scale bar, 5 µm (Horizontal arrow) and 50 s (Vertical arrow). **D)** Plot from (C) showing mean ± standard deviation of fold change of MAP7D1 intensity outside seeds with PTX or DDM normalized to before wash-in (PTX n = 16 microtubules, DDM n = 15 microtubules; from 3 experiments). A Welch’s t-test was used to compare between PTX and DDM, *** = p<0.001. **E)** Schematic depicting the model of how MAP7-family proteins mediate Kinesin-1 microtubule binding and selectivity. AlphaFold 3-predicted structures of KIF5B(1–560), alone and in complex with the MAP7D1 C-terminal region, are shown alongside the schematic. K560 requires MAP7-family proteins for release of autoinhibition and subsequent binding to microtubules. Activation by the diffuse MAP7D3 allows K560 to bind to all microtubules while MAP7D1 activates K560 specifically on stable microtubules. MAP7D1 itself confers this selectivity through binding to stable microtubules partially by recognizing detyrosination but primarily by recognition of the expanded lattice.

## DISCUSSION

Microtubules are organized into functionally distinct subsets that recruit specific motors and MAPs to support intracellular organization and transport. Although PTMs have long been proposed to define these subsets, the mechanisms establishing microtubule identity have remained incompletely understood. Here, we show that Kinesin-1 subset selectivity is not exclusively encoded within its motor domain and does not depend directly on microtubule PTMs. Instead, selective transport arises through MAP7D1, which recognizes expanded microtubule lattices and recruits Kinesin-1 to these microtubules through interactions with the Kinesin-1 CC1 region. Together, our findings identify microtubule lattice state as a major determinant of microtubule subset identity and selective motor recruitment.

Our results provide an explanation for the longstanding observation that Kinesin-1 preferentially moves over stable and highly modified microtubules in cells despite the relatively modest effects of tubulin PTMs on motor motility *in vitro* (Cai et al., 2009; Kaul et al., 2014; Reed et al., 2006; Sirajuddin et al., 2014). Previous work established MAP7 proteins as essential cofactors for Kinesin-1 activation and processivity (Barlan et al., 2013; Hooikaas et al., 2019; Monroy et al., 2018). We extend these findings by showing that different MAP7 family members confer distinct transport behaviors: MAP7D3 functions as a general activator that enables non-selective Kinesin-1 binding to microtubules, whereas MAP7D1 selectively recruits Kinesin-1 to stable microtubules that have expanded lattices. Thus, the microtubule preference of Kinesin-1 is not an intrinsic property of the motor itself but instead emerges from the differential localization of its activating cofactors. This could enable cells to tune Kinesin-1 selectivity by adjusting the relative expression levels of MAP7 family members, with MAP7D1 being the dominant activator in COS-7 and U2OS cells.

By generating a quadruple knockout cell line, we could demonstrate that acetylation and detyrosination are not required for MAP7D1 localization and Kinesin-1 selectivity, nor for forming a stable microtubule subset. These results do not imply that these PTMs are irrelevant for microtubule organization. Instead, our results suggest a hierarchical model in which lattice state acts upstream of PTMs. Stable microtubules acquire expanded lattices that are preferentially recognized by MAP7D1, and, as previously reported, by enzymes responsible for detyrosination and acetylation (Egoldt et al., 2025; Yue et al., 2023). As a result, PTMs and lattice state become tightly correlated in cells despite representing mechanistically distinct features of the microtubule polymer. In this model, PTMs could reinforce or consolidate subset identity rather than establish it, for example by promoting or hindering the binding of MAPs that impact microtubule stability (Chen et al., 2021; Lacroix et al., 2010; Peris et al., 2009; Szczesna et al., 2022; Valenstein & Roll-Mecak, 2016), or by protecting the microtubules from mechanical damage, as shown for acetylation (Portran et al., 2017; Xu et al., 2017). Consistently, we have observed that detyrosination positively impacts the binding of MAP7D1 to microtubules. This framework reconciles the strong correlation between PTMs and motor transport observed in cells with the relatively weak effects of individual PTMs on motor behavior *in vitro*.

Previous *in vitro* experiments demonstrated that binding of Kinesin-1 monomers in the nucleotide-free state can promote lattice expansion and stabilization (Peet et al., 2018), and suggested that motor binding can induce long-range conformational changes within the microtubule lattice (Muto et al., 2005; Shima et al., 2018). Furthermore, StableMARK, a dimeric rigor mutant of Kinesin-1 in which the motor domains are most likely mimicking the nucleotide-free, pre-powerstroke state (Rice et al., 1999), preferentially binds expanded lattices (de Jager et al., 2025; Jansen et al., 2023). In addition, motor-driven transport has been linked to lattice damage and subsequent repair of the microtubule polymer, sparking the idea that motors impact microtubule identity (Andreu-Carbo et al., 2024; Triclin et al., 2021). However, our observations suggest that, within cells, motor stepping alone is insufficient to alter microtubule identity. Upon overexpression of K370, the motor that moves indiscriminately over all microtubules, a subset of highly modified microtubules remains clearly visible (Fig. 1D). In these conditions, both StableMARK and MAP7D1 also retain their localization to a subset of stable and expanded microtubules (Fig. S1C, Fig. S3A). If motor movement were sufficient to drive widespread stabilization by either lattice expansion or damage and repair, one would instead expect all microtubules to become stabilized by the expression of motile motor domains that interact with all microtubules, resulting in redistribution of both acetylation and MAP7D1. Thus, our observations indicate that, while Kinesin-1 might have increased affinity for expanded lattices during specific steps of its mechanochemical cycle, the direct lattice remodeling of microtubules by walking motors occurs inefficiently inside the cell.

Nonetheless, recent work reported that strong overexpression of K560 reduces the level of acetylated microtubules (Andreu-Carbo et al., 2024). Given that K560, but not K370, engages MAP7-family proteins through its CC1 domain, this suggests that an abundance of Kinesin-1-MAP7 complexes can negatively impact microtubule stability. This could either reflect a lattice-destabilizing interplay between motile kinesins and immobile MAP7 or damage induced by buckling of microtubules as a result of microtubule-microtubule sliding caused by Kinesin-1-MAP7 pairs bound in *trans* to different microtubules. Indeed, recent work has demonstrated that force-induced microtubule buckling triggers microtubule damage beyond the limits of repair (Nandakumar et al., 2026).

The diffuse localization of K560 following depletion of MAP7D1 and MAP7D3 indicates that this commonly used truncated motor is not constitutively active in cells. This is consistent with the observation that, although K560 displays robust motility in purified systems, MAP7 family proteins dramatically enhance its motility in these conditions, even when they lack their microtubule binding domain (Hooikaas et al., 2019; Tan et al., 2023). Recent structural work has shown that the intrinsically disordered region between the Kinesin-1’s neck coil and CC1 enables the CC1 domain to fold back onto the motor domain in a conformation that would prevent motility (Ashaduzzaman et al., 2026; Tan et al., 2023). Consistently, our AlphaFold 3 modeling of K560 revealed a similar interaction between CC1 and Kinesin-1’s motor domain (Fig. 6E). Interestingly, when modeled in complex with the MAP7D1 C-terminus, this interaction was abolished and Kinesin-1 instead adopted an extended conformation where CC1 was aligned with the neck coil (Fig. 6E). We therefore propose that MAP7 binding interferes with the inhibitory interaction between CC1 and the motor domain, promoting an active kinesin-1 conformation.

The localization of MAP7D1 to the subset of microtubules with expanded lattices ensures Kinesin-1’s preferential motility over this subset, but how does MAP7D1 recognize expanded lattices? Structural studies indicate that the MAP7 microtubule-binding domain displays an extended interaction surface spanning multiple tubulin subunits and the tubulin C-terminal tails (Adler et al., 2024; Ferro et al., 2022), but unstructured regions not visible in cryo-electron microscopy are known to also contribute to microtubule binding through tubulin C-terminal tail interaction (Ferro et al., 2022; Hooikaas et al., 2019; Luo et al., 2020). Additionally, lattice expansion increases the accessibility of the tubulin C-terminal tails (Hotta et al., 2026), providing a potential basis for sensitivity to lattice conformation. Recent work also linked MAP7 recruitment to markers associated with lattice expansion during osmotic adaptation (Shen & Ori-McKenney, 2024). Our direct manipulation of lattice spacing now shows that expansion itself is sufficient to strongly promote MAP7D1 binding. However, the molecular basis by which MAP7D1 recognizes the expanded microtubule lattice remains to be determined.

In summary, our work demonstrates that microtubule lattice conformation is an important source of information within the cytoskeleton. In addition to tubulin nucleotide state and PTMs, microtubule lattice conformation constitutes a structural code that is interpreted by motors, MAPs, and modifying enzymes to generate functional diversity within the microtubule network. Determining how lattice states are established and recognized, and identifying additional readers and writers of the conformational states of the lattice will be important for understanding how the cytoskeleton organizes intracellular architecture and dynamics.

## ACKNOWLEDGEMENTS

We thank Toni van Capel for help with Fluorescence Assisted Cell Sorting, Hugo Snippert for ITPN constructs, J. Fernando Díaz for discodermolide and Anastasis Perrakis for VASH/SVBP. This work was supported by the Netherlands Organization for Scientific Research (NWO) via a VICI grant (project no. VI.C.212.062) to L.C.K. and an ENW-XL grant (project no. OCENW.XL21.XL21.048) to A.A. and L.C.K.

## AUTHOR CONTRIBUTIONS

Formal analysis (applying analytical techniques), J.v.S.; Investigation, J.v.S., V.M., M.K.I., D.J., N.P.; Methodology, W.N., J.v.S., V.M., M.K.I., D.J., N.P.; Resources, W.N., W.v.S., A.M., J. B.; Visualization (presenting data), J.v.S.; Writing – original draft, J.v.S., L.C.K.; Writing – review & editing, all authors; Funding acquisition, L.C.K. and A.A.; Supervision, L.C.K.;

## DECLARATION OF GENERATIVE AI AND AI-ASSISTED TECHNOLOGIES

During the preparation of this work, the author(s) used ChatGPT and Claude in order to check grammar and improve phrasing of the text. After using this tool or service, the authors reviewed and edited the content as needed and take full responsibility for the content of the publication.

## METHODS

### KEY RESOURCES TABLE

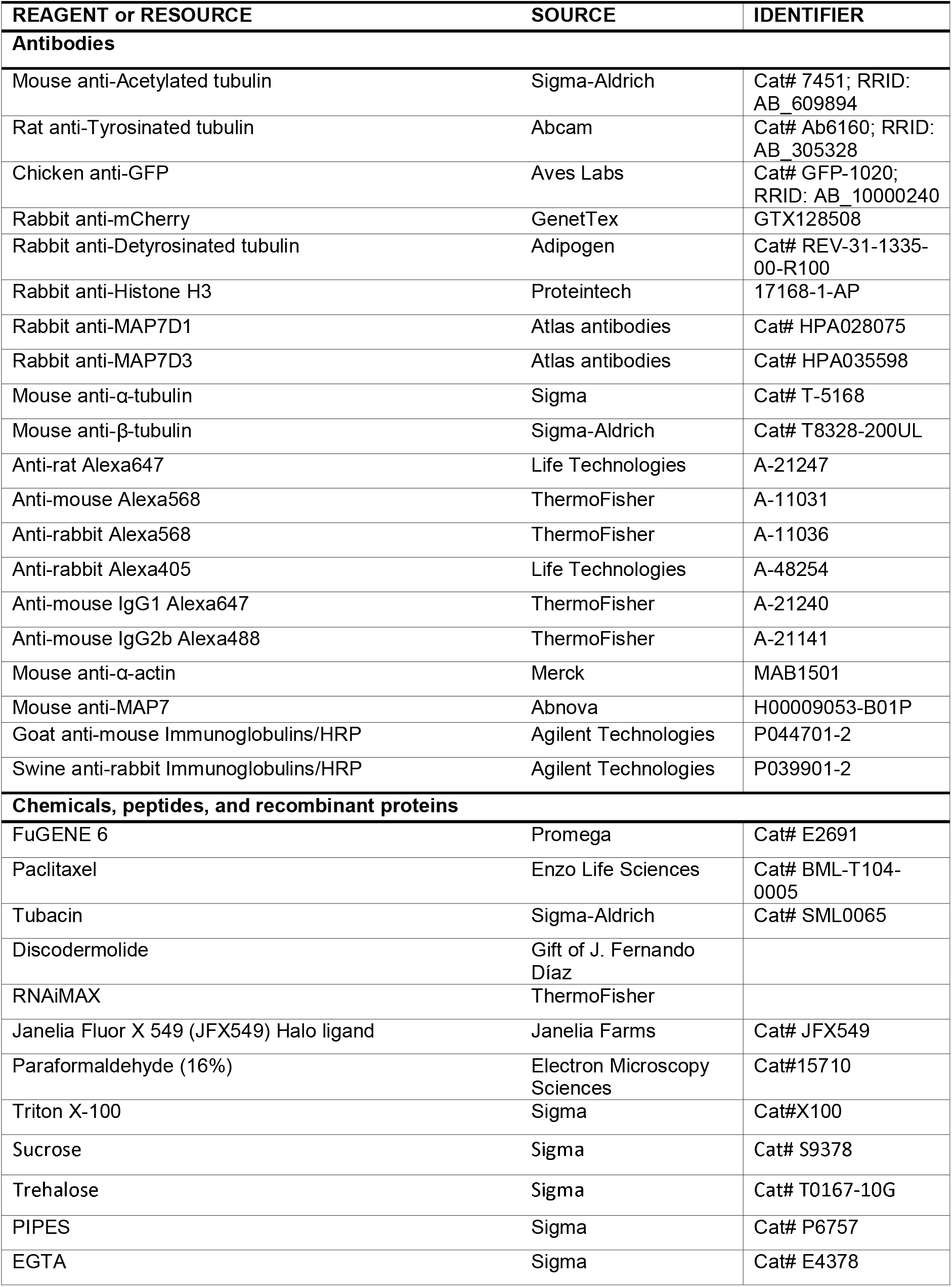
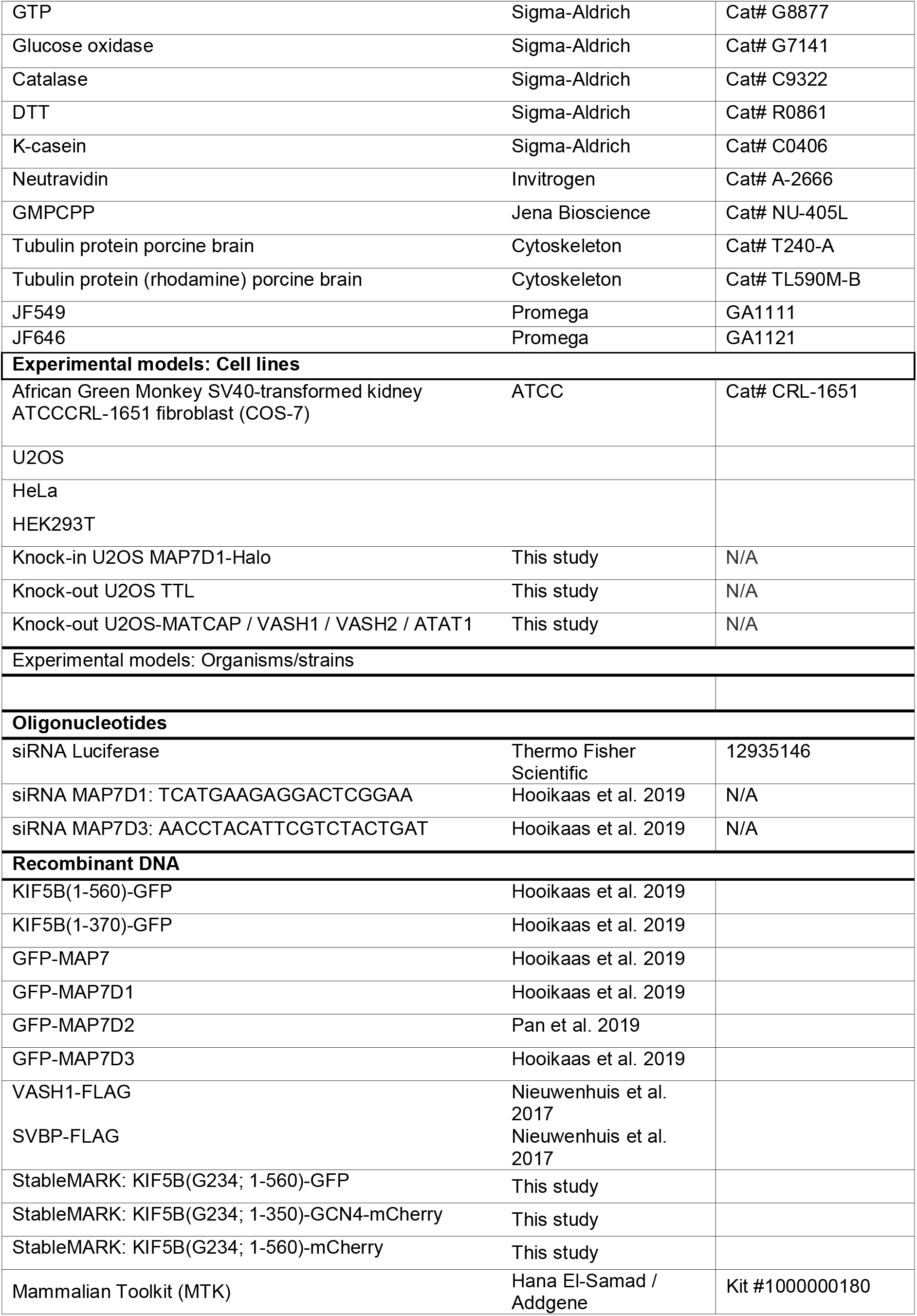
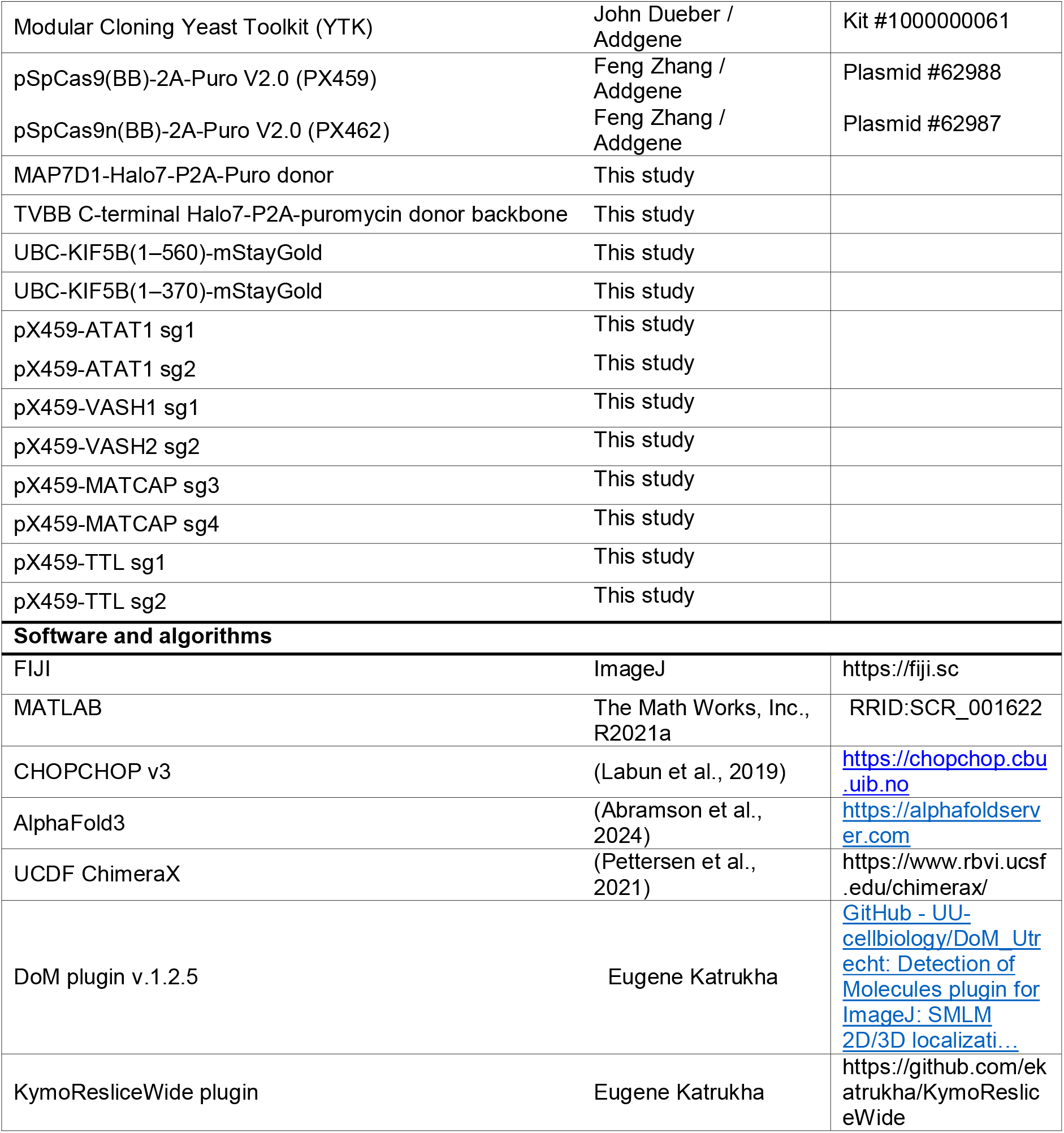

### Cell culture, transfection and knockdowns

COS-7 and U2OS cells were cultured in Dulbecco’s Modified Eagle Medium (DMEM; Capricorn Scientific) supplemented with 10% Fetal Bovine Serum (FBS; Corning) and 50 µg/mL penicillin/streptomycin at 37 °C with 5% CO2 and routinely tested for mycoplasma using the LT07-518 Mycoalert assay (Lonza). Cells were plated on 18-mm coverslips (immunolabeling experiments) or 25-mm coverslips (live-cell experiments) 1-2 days before transfection. For transient transfection, cells were transfected with FuGENE 6 (Promega) and DNA at a ratio of 3:1 following manufacturer’s protocol. For knockdowns, cells were transfected with 3µL lipofectamine RNAiMAX (ThermoFisher) and a total of 20 nM siRNA following manufacturer’s protocol. For combined knockdown and overexpression experiments, cells were transfected with siRNAs 1 day after plating followed by medium refreshment after 6-8h. At 32 h after siRNA transfection, cells were transfected with overexpression constructs. Finally, after a total incubation period of 52 h for siRNA and 20h for overexpression, cells were analyzed.

### Expression plasmids

To express truncated kinesin-1 motors tagged with mStayGold (Ivorra-Molla et al., 2024), expression plasmids encoding human KIF5B(1–560)-mStayGold or KIF5B(1–370)-mStayGold were assembled by one-step Golden Gate assembly using parts from the Mammalian Toolkit (MTK) and the Modular Cloning Yeast Toolkit (YTK) (Fonseca et al., 2019; Lee et al., 2015). The Modular Cloning Yeast Toolkit was a gift from John Dueber (Addgene kit #1000000061), and the Mammalian Toolkit was a gift from Hana El-Samad (Addgene kit #1000000180). The KIF5B motor and mStayGold were separated by a 24-amino-acid glycine-serine linker. Expression was driven by the human ubiquitin C (UBC) promoter. All assembled plasmids were verified by whole-plasmid sequencing (Plasmidsaurus). Construct details are provided in the Key Resource Table.

### Generation of knockout cell lines

U2OS knockout cell lines were generated by CRISPR-Cas9 genome editing. Guide RNAs were designed using CHOPCHOP (Labun et al., 2019), cloned into pSpCas9(BB)-2A-Puro (PX459) V2.0 (Ran et al., 2013);(Addgene plasmid #62988), and transiently transfected into U2OS cells. Cells were subjected to two consecutive rounds of transfection separated by transient puromycin selection and recovery. Single-cell clones were isolated by limiting dilution and screened by immunofluorescence. Guide RNA sequences are provided in Table S1.

**ATAT1 knockout:** An **ATAT1** knockout U2OS cell line (clone B7) was generated using guide RNAs targeting exon 6 and the exon 10 splice donor. Clones were screened for loss of acetylated α-tubulin by immunofluorescence. Clone B7 was confirmed by targeted amplicon sequencing (Plasmidsaurus) to contain only knockout alleles for ATAT1.

**TTL knockout:** A **TTL** knockout U2OS cell line (clone F2) was generated using the same workflow. Clones were screened by immunofluorescence for loss of tyrosinated α-tubulin and a corresponding increase in detyrosinated α-tubulin. Clone F2 was used for all subsequent experiments.

**VASH1/VASH2/MATCAP/ATAT1 quadruple knockout:** A quadruple knockout U2OS cell line was generated by sequential CRISPR-Cas9 genome editing starting from the validated ATAT1 knockout clone B7. VASH1 and VASH2 were disrupted in two consecutive rounds of genome editing, after which MATCAP was targeted using two guide RNAs. Single-cell clones were isolated by limiting dilution and screened by immunofluorescence for loss of detyrosinated α-tubulin. Clone B-H8 was confirmed by targeted amplicon sequencing (Plasmidsaurus) to contain only knockout alleles for VASH1, VASH2, MATCAP, and ATAT1.

### Generation of knock-in cell lines

MAP7D1-Halo7 polyclonal knockin U2OS lines were generated by endogenous tagging using in-trans paired nicking (ITPN), in which Cas9 D10A nickase simultaneously nicks the genomic target locus and donor plasmid to promote homology-directed repair without generating genomic double-strand breaks. Guide RNAs were designed to target sequences overlapping the endogenous stop codon, enabling C-terminal tagging.

Guide RNA expression plasmids were generated by cloning guide sequences into pSpCas9n(BB)-2A-Puro V2.0 (PX462; Addgene #62987). Gene-specific donor plasmids were generated by SapI Golden Gate assembly of 500-600 bp homology arms into ITPN donor backbones described in (Bollen et al., 2022). The donor backbones encoded GGGGSAS-Halo7-P2A-puromycin repair cassettes immediately upstream of the endogenous stop codon. A TVBB C-terminal mNeonGreen-P2A-puromycin donor backbone was a gift from Hugo Snippert (Addgene plasmid #169228 (Bollen et al., 2022)). The corresponding TVBB Halo7-P2A-puromycin donor backbone was generated for this study by replacing the mNeonGreen cassette with a synthetic Halo7-P2A-puromycin cassette using Gibson assembly. Guide plasmids were verified by Sanger sequencing, and donor plasmids were verified by whole-plasmid sequencing (Plasmidsaurus).

U2OS cells were co-transfected with guide RNA expression plasmids and donor plasmids using Fugene6 transfection reagent (Promega). Transient puromycin selection was used to enrich for transfected cells. Following recovery from transient puromycin selection, Halo7-positive cells were enriched by fluorescence-activated cell sorting (FACS). Halo7-expressing cells were labeled with JF646 HaloTag ligand prior to sorting. MAP7D1-Halo7 cell lines were maintained as polyclonal populations following fluorescence enrichment. Fluorescent populations were selected based on fluorescence intensity and subcellular localization consistent with endogenous MAP7 protein distribution.

### Drug treatments

Cells were incubated with 2 µM paclitaxel (Enzo Life Sciences BML-T104-0005) or 200 nM discodermolide for 20 min, and 10 µM Tubacin for 2 hr. Drugs were diluted in fresh, prewarmed medium and well-mixed prior to addition. For U2OS KO validation experiments, cells were treated with 2µM paclitaxel for 3 hr before lysis.

### Immunofluorescence

For staining of endogenous MAP7 proteins, cells were fixed at 37 °C with prewarmed fixation buffer (4% v/v PFA (Electron Microscopy Sciences 15,710) and 4% w/v sucrose in MRB80) for 12min. To visualize overexpressed motor proteins bound to microtubules, an extraction fixation protocol was optimized and used for all experiments with immunofluorescence after overexpression. First cells were extracted at 37 °C for 30-45 s with prewarmed extraction buffer (0.3% v/v Triton X-100 (Sigma X100) and 0.1% v/v glutaraldehyde (Sigma G7526) in MRB80 then fixed for 12min at 37 °C with fixation buffer (4% v/v PFA and 4% w/v sucrose in MRB80). For fixations with glutaraldehyde, cells were quenched three times for 5 min with quenching buffer (5 mg/ml sodium borohydride in PBS). Cells were washed 2 times with wash buffer (0.1% Triton X-100 in PBS) and incubated with blocking buffer (3% w/v BSA, 0.1% v/v Triton X-100 in PBS) for 30 min. Cells were incubated overnight at 4 °C with primary antibodies diluted in blocking buffer, washed 3 times with wash buffer and incubated with secondary antibodies diluted in blocking buffer for 2 h at room temperature. Cells were washed 3 times with wash buffer, 1 time with Milli-Q, dipped in 96% ethanol and dried. Coverslips were mounted using ProLong Diamond antifade mounting medium.

### Fixed-cell imaging

Images of fixed cells were acquired using a Zeiss LSM980 confocal microscope equipped with Airyscan 2 on an Axio Observer 7 inverted microscope stand (Carl Zeiss). Images were acquired in Airyscan super-resolution (SR) mode using an Alpha Plan-Apochromat 100×/1.46 NA oil-immersion objective and laser excitation at 405, 445, 488, 561, and 639 nm. Fluorescence was detected using the Airyscan detector, and images were reconstructed using the Airyscan processing algorithm in ZEN Blue software (version 3.9, Carl Zeiss).

### Image processing and analysis

FIJI (version 2.14.0) was used to analyze microscopy data (Schindelin et al., 2012). Matlab was used to visualize data output and for statistical analysis. To quantify colocalization between microtubule markers, 10 × 10 µm regions of interest (ROIs) were manually selected in the peripheral cytoplasm, where individual microtubules could be resolved with minimal overlap. Pearson’s correlation coefficients between fluorescence channels were calculated for each ROI using the JaCoP plugin in FIJI (Bolte & Cordelieres, 2006). For each cell, one or more ROIs were analyzed as indicated in the figure legends. Individual ROIs were treated as independent measurements and grouped by biological replicate for visualization. Statistical analyses were performed in MATLAB using a Wilcoxon matched-pairs signed-rank test for paired comparisons within the same ROIs and Welch’s t-test for comparisons between experimental conditions, as specified in the figure legends.

### Protein sequence alignment and conservation analysis

Protein sequences were aligned in MATLAB using the Bioinformatics Toolbox multialign function with the BLOSUM62 substitution matrix and affine gap penalties (opening: 10; extension: 0.5). Conservation was calculated as the fraction of sequences containing the most frequent amino acid at each position, including gaps in the total sequence count, and visualized using a cyan-to-orange color scale.

### Western blot

For Western blot analysis, HeLa, COS-7, U2OS, and HEK cells were washed with 1X PBS and lysed in Laemmli buffer (4% SDS, 20% glycerol, 120 mM Tris pH 6.8). The Pierce Rapid Gold BCA Protein Assay Kit (Thermo, A53226) was used to determine total protein concentration. 20 µg of all cell lines was loaded onto a 4-15% precast polyacrylamide gel (Biorad, 4568084). After protein separation during SDS-PAGE, proteins were transferred to a nitrocellulose membrane for immunoblotting. The transfer was verified using 0.1% Ponceau (ChemCruz, C2422). Subsequently, the membrane was blocked in 1X PBS + 3% BSA (Carl Roth, 8076.3) and incubated overnight at 4 ⁰C using the following primary antibodies: mouse anti-MAP7 (Abnova, H00009053-B01P, 1:2000), rabbit anti-MAP7D1 (Atlas, HPA028075, 1:1000), rabbit anti-MAP7D3 (Atlas, HPA035598, 1:1000), and anti-α-actin (Merck, MAN1501, 1:10.000). The membrane was then washed with TBST (0.1% Tween20 in Tris-buffered saline) and incubated for 1 hr at room temperature using HRP conjugated secondaries. The following secondary antibodies were used: Goat anti-mouse Immunoglobulins/HRP (Agilent Technologies, P044701-2, 1:2000), and Swine anti-rabbit Immunoglobulins/HRP (Agilent Technologies, P039901-2, 1:2000). After washing, ECL Western Blotting Substrate (Promega, W1015) was added to visualize the protein bands and incubated for 2 min. Finally, membranes were scanned on an Amersham ImageQuant 800 imaging system.

### Purification of MAP7D1 for in-vitro reconstitution assays

MAP7D1 was purified from HEK293T cells using the Strep(II)-strepactin affinity purification as described previously (Meiring et al., 2026). Briefly, cells were transfected with a 1:3 w/w mix of the construct and polyethyleneimine (Polysciences). The cells were harvested 26h after transfection in ice-cold lysis buffer (50mM HEPES, 1 mM MgCl_2_, 1 mM DTT, 0.1mM ATP, 300 mM NaCl, and 0.5% Triton X-100; pH 7.4) containing EDTA free protease inhibitors (Roche). They were incubated on ice for 15 mins, centrifuged and the supernatant was incubated in Streptactin Sepharose beads (IBA lifesciences) for 1hr. The beads were washed 3 times with lysis buffer but with 150mM NaCl instead of 300mM (low salt washes) and three times with lysis buffer with 300mM NaCl (high salt washes). The protein was then eluted in elution buffer (50 mM HEPES, 150 mM NaCl, 1 mM MgCl_2_, 1 mM EGTA, 1 mM DTT, 0.05% Triton X-100 and 2.5 mM d-desthiobiotin (Sigma-Aldrich), pH 7.4). The proteins were then aliquoted, snap frozen and stored at -80°C.

VASH/SVBP were co-expressed in Spodoptera frugiperda (Sf9) insect cells using the Invitrogen Bac-to-Bac method as described previously (Nieuwenhuis et al., 2017). The complex was expressed in Sf9 insect cells using Insect-Xpress media (LONZA). Cells were grown in a shaker at 28 °C and infected at a density of 2×106 cellsml−1. Cells were harvested 72h post-infection, and the biomass was flash frozen in liquid nitrogen and stored at −20 °C. The purification protocol described previously (Nieuwenhuis et al., 2017) was used with the following modifications. The VASH1–SVBP complex was affinity purified using Streptactin superflow resin (IBA) using a 20mM HEPES/NaOH pH 7.5, 500mM NaCl, 1mM Tris(2-carboxyethyl) phosphine, and 5 mM Desthiobiotin elution buffer. To further purify the complex, size-exclusion chromatography was employed, using a Superdex S-200 10 300 column (GE Healthcare) in 20mM HEPES/NaOH pH 7.5, 150mM NaCl and 1mM Tris(2-carboxyethyl) phosphine. The fractions containing the purified protein were pooled and subsequently concentrated with a 10 kDa centrifugal concentrator (Amicon), flash frozen and stored at −80 °C until further use.

### *In vitro* reconstitution assays

The flow chambers were prepared by plasma cleaning coverslips and using two strips of double-sided tape to create ∼10 µL volume chambers on microscope slides. The surface was functionalized by sequential 5-minute incubations with 0.2 mg/mL PLL-PEG-Biotin (Susos AG) and 0.83 mg/mL neutravidin (Invitrogen), both dissolved in MRB80 buffer (80 mM K-Pipes, 1 mM EGTA, 4 mM MgCl₂; pH 6.80 with KOH). Double-cycled biotinylated GMPCPP seeds prepared as described previously (Jansen et al., 2023) were then flowed in and incubated for 2 minutes, followed by a >3-minute incubation with κ-casein.

All tubulin reagents used were from Cytoskeleton Inc. All mixes contained a master mix composed of 0.1% [wt/vol] methylcellulose, 0.5 mg/ml K-casein, 50mM KCL, 50 mM glucose, 0.2 mg/ml catalase, 0.5 mg/ml glucose oxidase, and 10 mM DTT, and were mixed in the MRB80 buffer.

For the dynamic paclitaxel/discodermolide flow-in assays, the microtubule mix had master mix supplemented with 17 µM porcine brain tubulin, 0.6 µM TRITC rhodamine labelled tubulin and 1 mM GTP; and the wash-in mix had master mix with 20 µM paclitaxel or 200 nM discodermolide. For the experiments with MAP7D1 and taxol or discodermolide, the microtubule mix was supplemented with 17 µM porcine brain tubulin, 0.6 µM TRITC rhodamine labelled tubulin, 1 mM GTP and 500pM MAP7D1; and the wash in mix had 4.85 µM porcine brain tubulin, 0.15 µM TRITC rhodamine labelled tubulin, 20 µM paclitaxel or 200 nM discodermolide and 500pM MAP7D1. All mixes were spun in an Airfuge for 5 min at 119,000 x g. The microtubule mixes were flowed into the chambers, and the wash-in mixes were kept on ice before manually washing-in during image acquisition.

The immunofluorescence experiments were done on GMPCPP seeds. The seeds were first incubated for 45 mins with master mix supplemented with MRB80 or VASH/SVBP. A mix was made containing the master mix and MAP7D1, split, and flowed into both the chambers. After 10 mins, the seeds were fixed by flowing in a mix of 4% paraformaldehyde and 0.1% glutaraldehyde in MRB80. After three washes with 0.4% sodium borohydride in PBS the seeds were blocked with 3% BSA and 0.1% triton-X for 3 mins before incubating with the primary antibodies for 10 mins. They were then washed three times with 0.1% triton-X in PBS before a 5 min incubation in secondary antibodies and then washed 3 times with 0.1% triton-X in PBS.

### TIRF microscopy

Total internal reflection fluorescence (TIRF) microscopy was performed on an inverted Nikon Eclipse Ti-E research microscope (Nikon) equipped with azimuthal spinning TIRF illumination, a perfect focus system, and an ASI motorized MS-2000-XY stage. Imaging was performed using a Nikon APO TIRF 100× 1.49 NA oil immersion objective in combination with an iLas2 illumination system (Roper Scientific, now Gataca Systems). For *in vitro* assays, images were acquired with a CoolSNAP MYO CCD camera (Teledyne Photometrics) providing a pixel size of 0.045 μm/pixel. For live-cell experiments, images were acquired with an Evolve Delta 512 EMCCD camera (Photometrics) providing a pixel size of 0.064 μm/pixel. For simultaneous imaging of green and red fluorescence an ET-GFP/Cherry filter cube was used (Chroma, 59022), together with an Optosplit III (Cairn Research Ltd) equipped with double-emission filter cube configured with ET525/50m, ET630/75m, and T585lprx (Chroma). Samples were maintained at 30 °C for *in vitro* assays and 37 C with 5% CO2 for live-cell experiments using a Tokai Hit STXG-PLAMX-SETZ21L Stage Top Incubator. Excitation was achieved using the following lasers: Coherent OBIS 561 nm (100 mW), Vortran Stradus 642 nm (110 mW), and Vortran Stradus 488 nm (150 mW). Emission was collected through the corresponding Chroma filters ET-GFP (49002), ET-mCherry (49008), and ET-647. Image acquisition was controlled using Metamorph software (version 7.10.2.240, Molecular Devices).

### Image processing of *in vitro* assays

To calculate lattice expansion, a straight line was drawn from the end of the bright seed to a speckle toward the plus end of the kymograph before the wash-in. After the flow-in, a second line was drawn between the same two points. The difference in lengths, reflecting the speckle shift, was used to calculate the percentage of expansion. To calculate the intensity, the movies were averaged over frames, and the mean intensity was calculated at the required region of interest after background correction. The ImageJ plugin KymoResliceWide v.04 was used to generate Kymographs (https://github.com/ekatrukha/KymoResliceWide).

### Alpha-fold modeling

Protein structure prediction was performed using AlphaFold3 (Abramson et al., 2024) with two copies of KIF5B (amino acids 1–560) and two copies of MAP7D1 (amino acids 581–724). The predicted protein complex was subsequently visualized and color-coded using UCSF ChimeraX (Pettersen et al., 2021).

## SUPPLEMENTARY FIGURES

**Figure S1:**
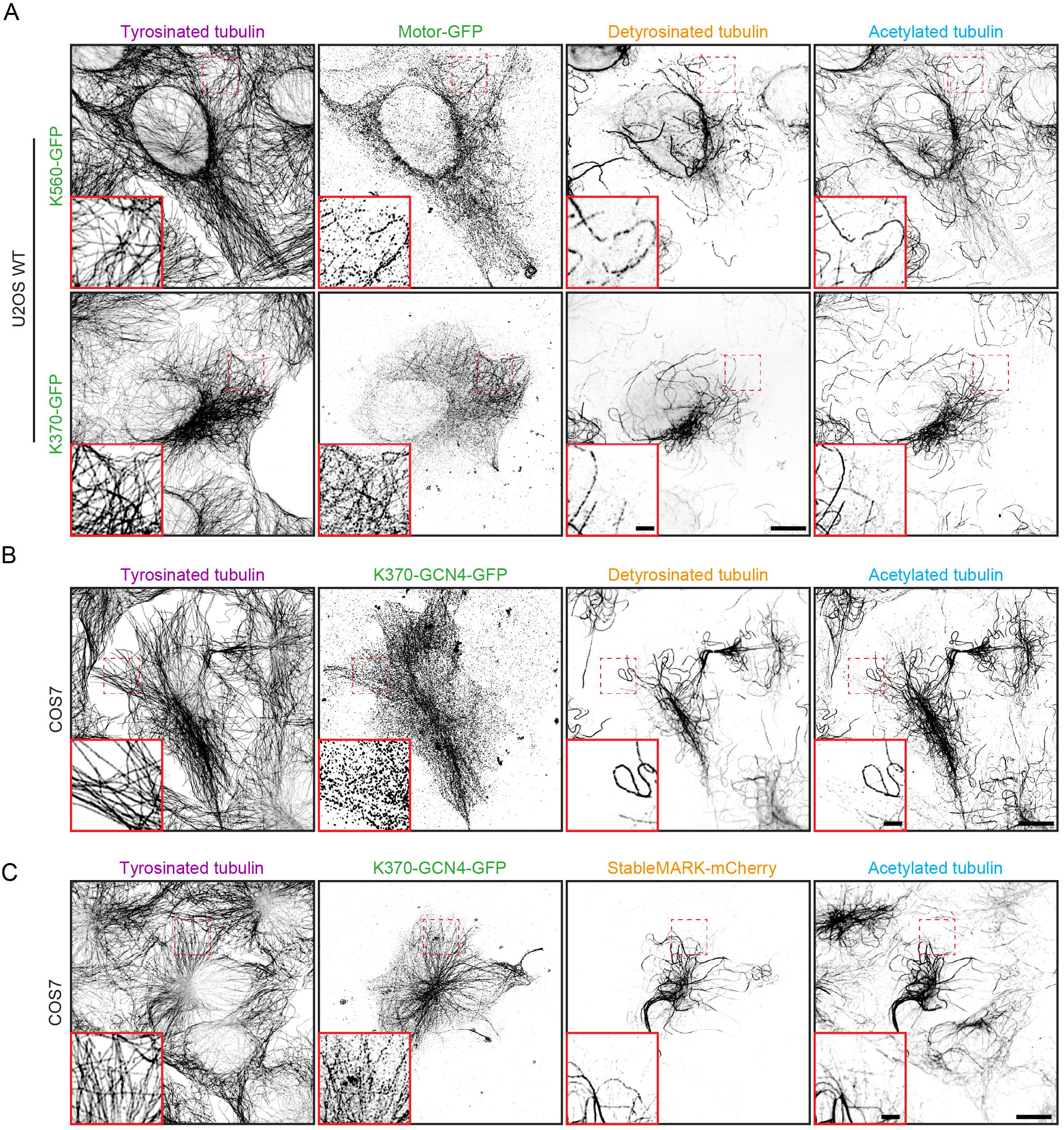
Kinesin-1 subset selectivity is conserved in U2OS cells and is not specified by the motor domain. **Related to Figure 1. A-B)** U2OS cells transfected at low levels with K560-GFP or K370-GFP (A) and COS-7 cells transfected at low levels with K370-GCN4-GFP (B), were fixed using an optimized extraction-fixation protocol to visualize microtubule-bound motor before staining for tyrosinated α-tubulin (Tyr), GFP, detyrosinated α-tubulin (Detyr) and acetylated α-tubulin at position K40 (Ace). **C)** COS-7 cells expressing K370-GCN4-GFP and StableMARK-mCherry were fixed and stained for tyrosinated α-tubulin (Tyr), acetylated α-tubulin at position K40 (Ace), GFP and mCherry. Scale bar, 10 µm (for whole cell) and 2 µm (for zoom).

**Figure S2:**
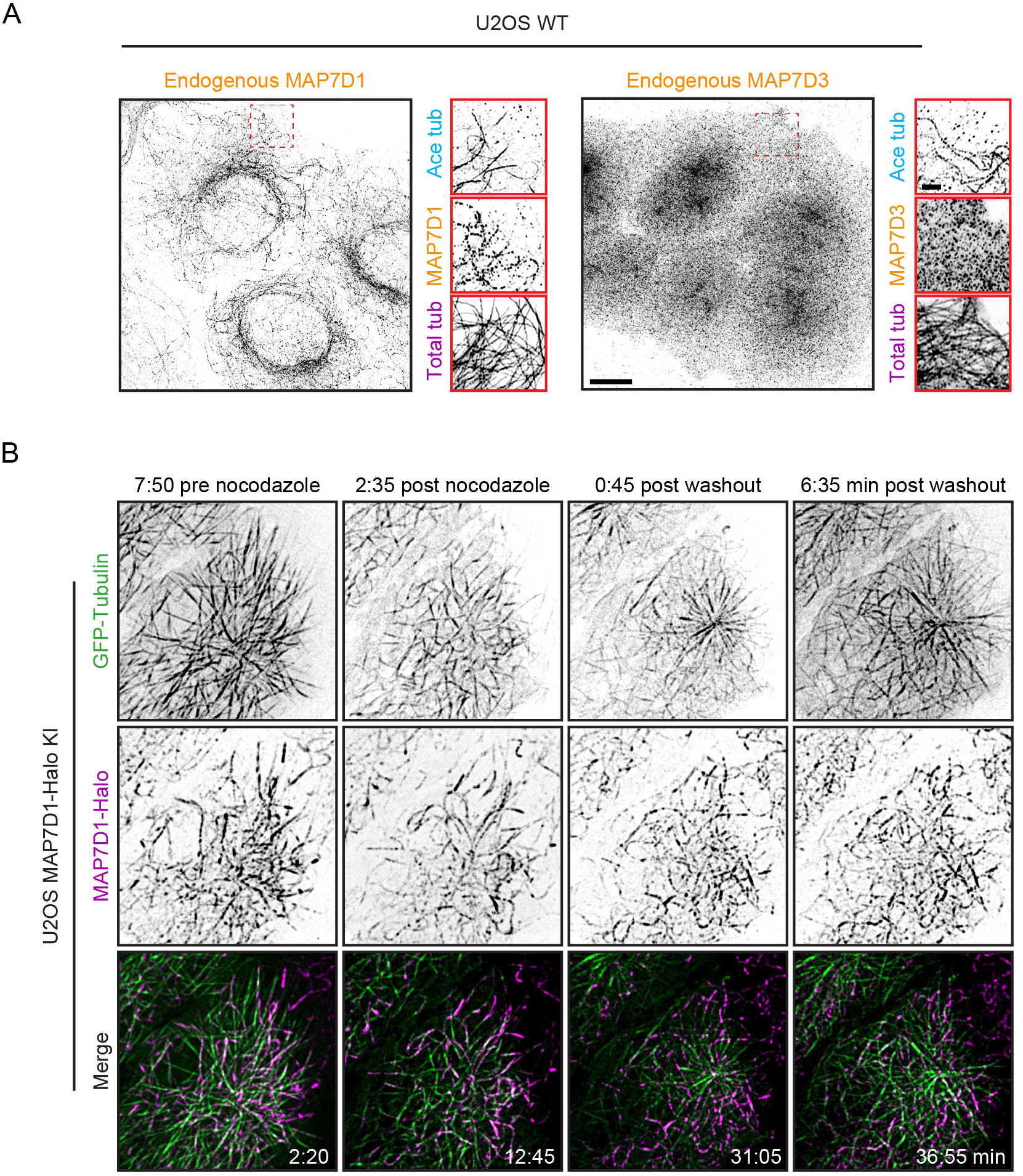
MAP7-family expression and localization to stable microtubules. **Related to Figure 2. A)** U2OS cells were fixed and stained for α-tubulin and β-tubulin (Total tub), acetylated α-tubulin at position K40 (Ace tub) and endogenous MAP7D1 or MAP7D3. **B)** U2OS MAP7D1-Halo knock-in cells expressing GFP-α-tubulin were labeled with JF549 HaloTag ligand and imaged using live-cell TIRF microscopy. Average intensity projections of five frames spanning 20 s, show microtubules before and during nocodazole treatment and following washout. MAP7D1-positive microtubules persist during nocodazole treatment, whereas newly polymerized microtubules following washout are MAP7D1-negative. Scale bars, 10 µm (A whole-cell images), 2 µm (A zooms) and 5µm (B).

**Figure S3:**
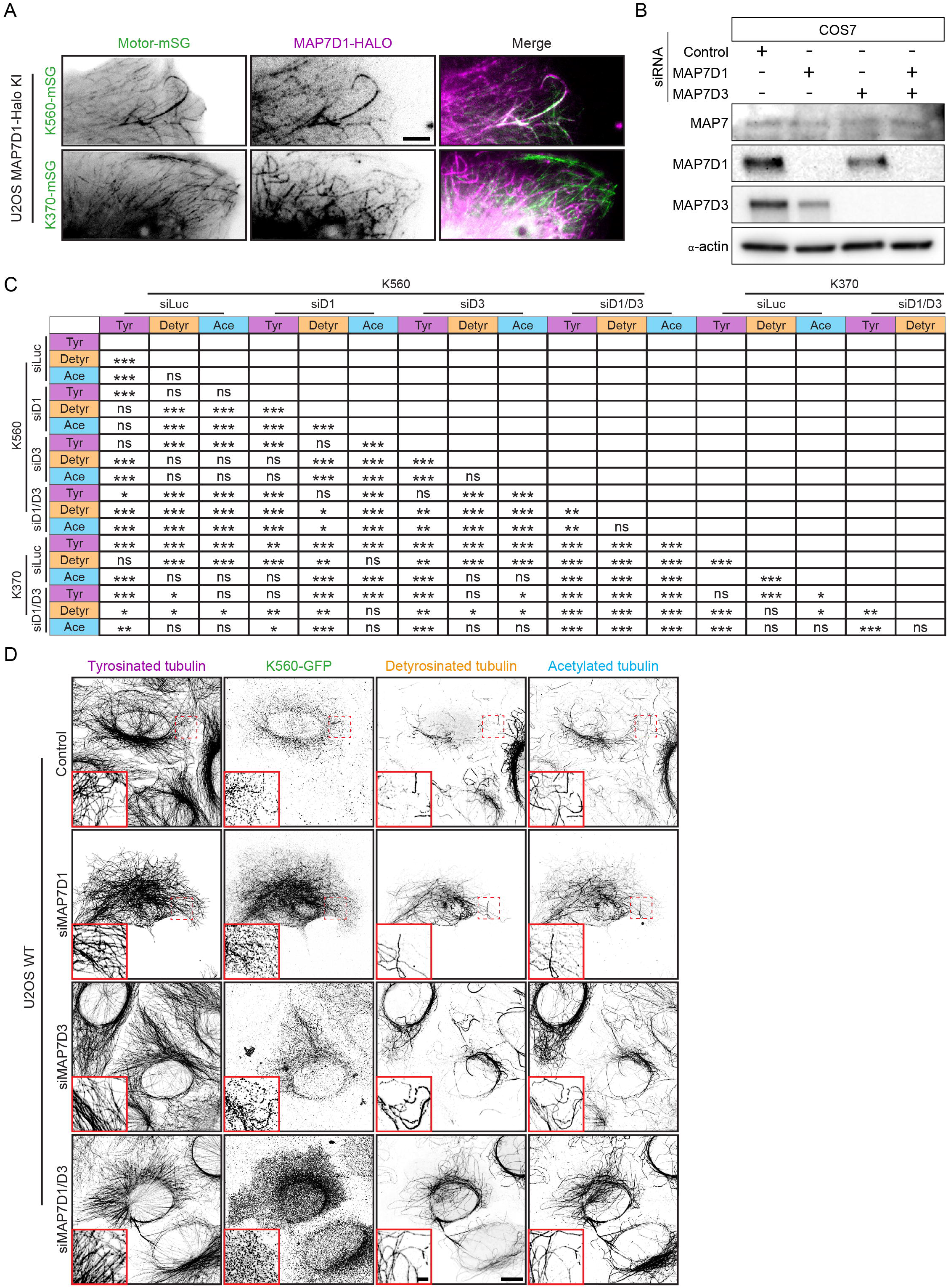
MAP7D1 determines Kinesin-1 microtubule subset selectivity. **Related to Figure 3. A)** U2OS MAP7D1-HALO knock-in cells expressing K560-mSG or K370-mSG were labeled with JF549 fluorescent HaloTag ligand and imaged using live-cell TIRF microscopy. Maximum projections of 20 frames covering 2 seconds show the preferential movement of K560 along MAP7D1-positive microtubules, whereas K370 also moves along MAP7D1-negative microtubules. **B)** Western blot analysis of COS-7 cells transfected with siRNAs targeting Luciferase (Control), MAP7D1, MAP7D3, or both MAP7D1 and MAP7D3. Blots were stained for the indicated MAP7-family proteins and β-actin to assess knockdown efficiency and expression of other MAP7-family members. **C)** Table showing statistical comparison of Pearson’s correlation coefficient between K560-GFP or K370-GFP and tyrosinated α-tubulin (Tyr), detyrosinated α-tubulin (Detyr) and acetylated α-tubulin at position K40 (Ace) for all siRNA conditions quantified in Fig. 3C. A Wilcoxon matched-pairs signed-rank test was used to compare correlations between microtubule subsets within the same condition, and a Welch’s t-test was used to compare correlations between conditions. **D)** U2OS cells transfected with siRNAs targeting Luciferase (Control), MAP7D1, MAP7D3 or both MAP7D1 and MAP7D3 were subsequently transfected at low levels with K560-GFP. Cells were fixed using an extraction-fixation protocol before staining for tyrosinated α-tubulin (Tyr), GFP, detyrosinated α-tubulin (Detyr) and acetylated α-tubulin at position K40 (Ace). Scale bars, 5 µm (A), 10 µm (D whole cell) and 2 µm (D zoom).

**Figure S4:**
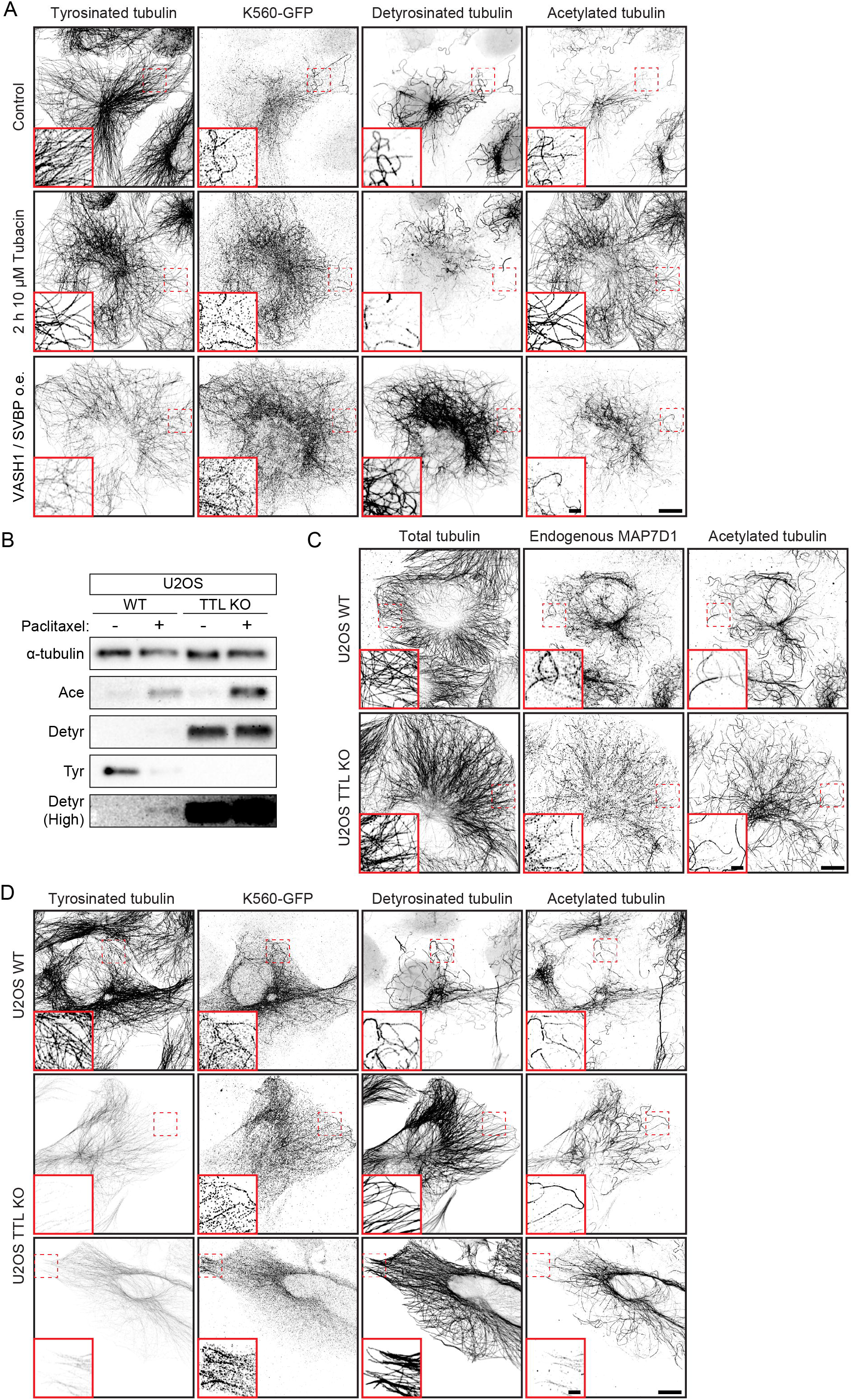
Effects of acetylation and detyrosination on MAP7D1 and Kinesin-1 localization. **Related to Figure 4. A)** COS-7 cells transfected at low levels with K560-GFP were treated with DMSO (control), 10µM Tubacin for 2 h or co-transfected with VASH1-FLAG/SVBP-FLAG before fixing and staining for tyrosinated α-tubulin, GFP, detyrosinated α-tubulin and acetylated α-tubulin at position K40. **B)** Western blot of wildtype U2OS and TTL KO U2OS cell extracts stained for α-tubulin, tyrosinated α-tubulin (Tyr), detyrosinated α-tubulin (Detyr) and acetylated α-tubulin at position K40 (Ace). **C)** Wildtype and TTL knockout U2OS cells were fixed and stained for endogenous MAP7D1, acetylated α-tubulin at position K40 and α-tubulin and β-tubulin (Total tubulin). **D)** Wildtype and TTL knockout U2OS cells transfected at low levels with K560-GFP were fixed and stained for tyrosinated α-tubulin, GFP, detyrosinated α-tubulin and acetylated α-tubulin at position K40. Scale bars, 10 µm (whole cells) and 2 µm (D zooms).

**Figure S5:**
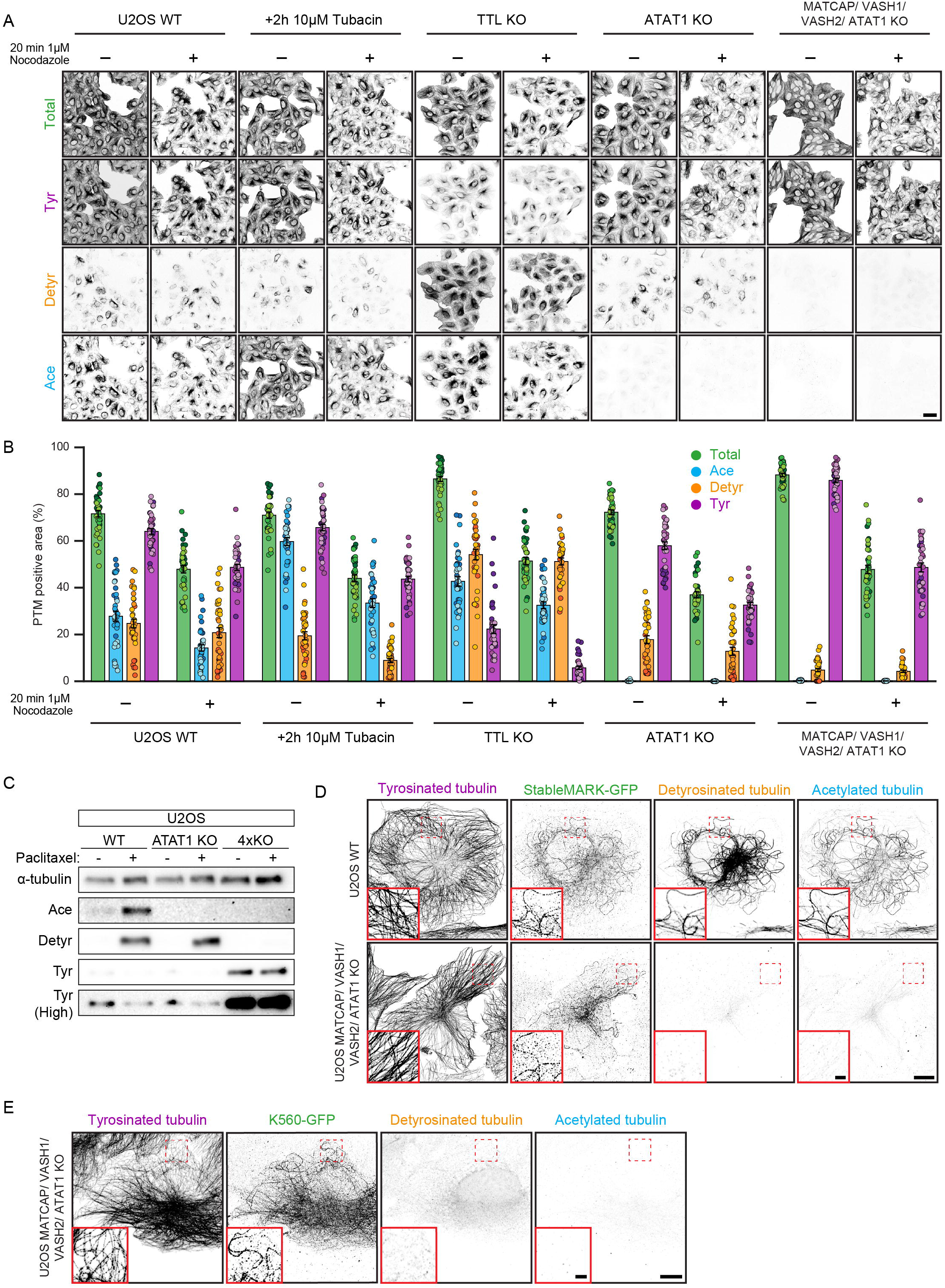
Stable microtubules persist in the absence of detyrosination and acetylation. **Related to Figure 5. A)** Wildtype, Wildtype treated with 10 µM Tubacin for 2 h, TTL KO, ATAT1 KO and MATCAP/VASH1/VASH2/ATAT1 quadruple knockout (4xKO) U2OS cells were treated with DMSO (Control) or 1 µM nocodazole for 20 min before fixation and staining for α-tubulin and β-tubulin (Total tubulin), tyrosinated α-tubulin (Tyr), detyrosinated α-tubulin (Detyr) and acetylated α-tubulin at position K40 (Ace). Representative lower-magnification images were used to quantify the percentage of cell area occupied by microtubules in Fig. 5D. **B)** Quantifications of (A). The percentage of cell area occupied by total tubulin signal was measured. Dots indicate individual cells and shades indicate experimental repeats (n = 39 cells from 3 independent experiments). **C)** Western blot of wildtype U2OS, ATAT1 KO and MATCAP/VASH1/VASH2/ATAT1 quadruple knockout (4xKO) U2OS cell extracts stained for α-tubulin, tyrosinated α-tubulin, detyrosinated α-tubulin and acetylated α-tubulin at position K40. **D-E)** Wildtype (D) and 4xKO (D,E) U2OS cells expressing low levels of StableMARK-GFP (D) or K560-GFP (E) were fixed and stained for tyrosinated α-tubulin, GFP, detyrosinated α-tubulin and acetylated α-tubulin at position K40. Scale bars, 50 µm (A), 10 µm (D,E whole cell) and 2 µm (D,E zoom).

**Figure S6.**
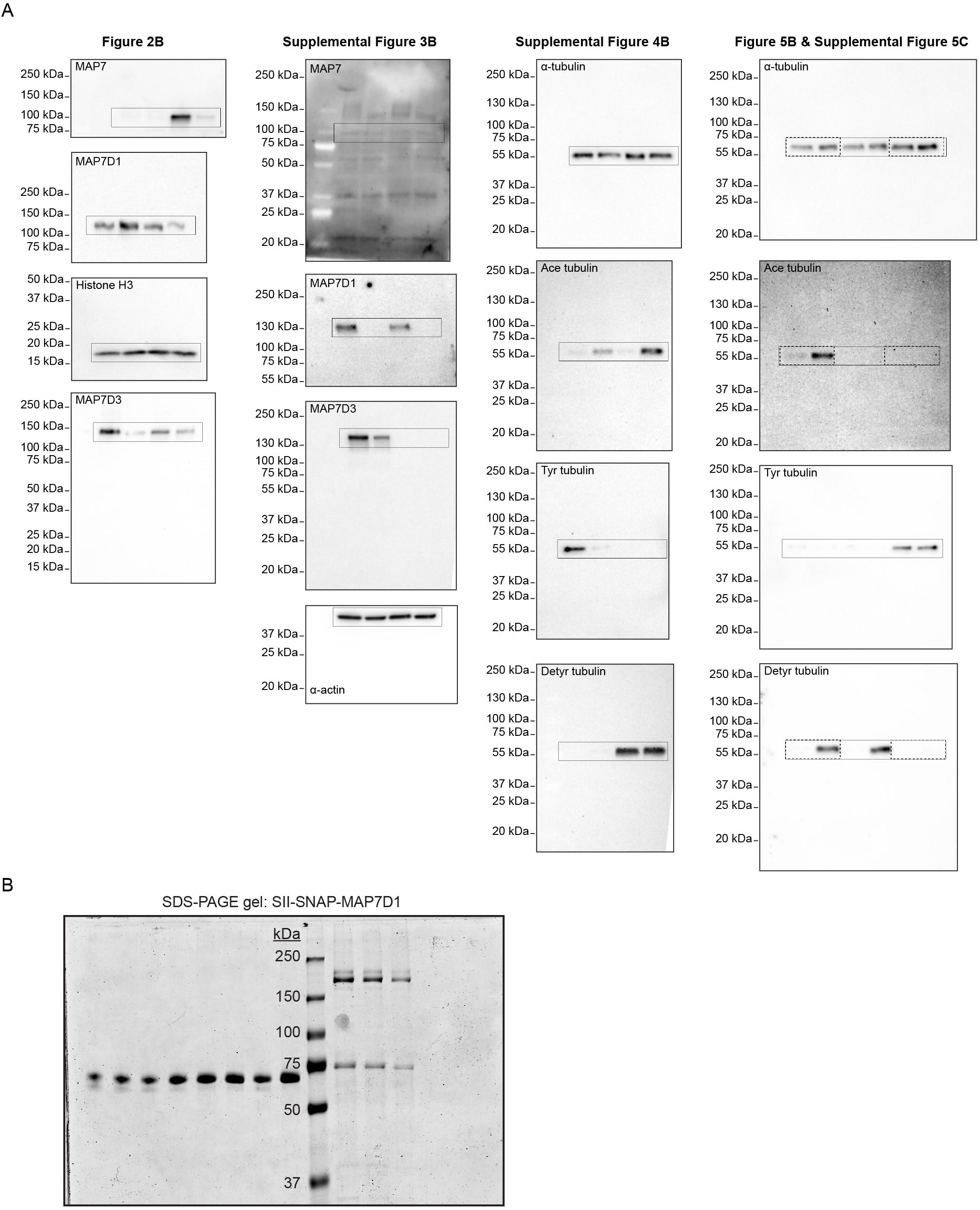
Full Western blots and SDS-PAGE gel. **(A)** Uncropped Western blots corresponding to the indicated figures. Solid rectangles indicate the regions shown in the respective figures, as labeled above each blot. For the rightmost blots, dashed rectangles indicate the regions shown in Fig. 5B, whereas solid rectangles indicate those shown in Fig. S5C. **(B)** SDS-PAGE gel showing the purification of SNAP-MAP7D1 used in Fig. 6C,D.

